## Supplement for "Community structure and temporal dynamics of SARS-CoV-2 epistatic network allow for early detection of emerging variants with altered phenotypes"

### Appendix A Supplementary information

#### A.1 Supplementary tables

| | Complete | 1st truncated | 2nd truncated |
|---|---|---|---|
| VOCs/VOIs | | | |
| Non-negative lag & significant ($p < 0.05$, Student's $t$-test) medium-to strong negative correlation for VOCs/VOIs across all countries | 70% | 74% | 77% |
| CI $\rho$ | $[-0.97, 0.39]$ | $[-0.97, -0.36]$ | $[-0.97, -0.44]$ |
| Mean $\rho$ | -0.73 | -0.74 | -0.74 |
| CI $l^*$, days | $[0, 168]$ | $[0, 168]$ | $[0, 140]$ |
| Mean $l^*$, days | 23.5 | 20.5 | 20.6 |
| VOCs only | | | |
| Non-negative lag & significant ($p < 0.05$, Student's $t$-test) medium-to strong negative correlation for VOCs across all countries | 76% | 84% | 89% |
| CI $\rho$ | $[-0.91, -0.36]$ | $[-0.95, -0.36]$ | $[-0.97, -0.37]$ |
| Mean $\rho$ | -0.74 | -0.72 | -0.72 |
| CI $l^*$, days | $[0, 140]$ | $[0, 168]$ | $[0, 168]$ |
| Mean $l^*$, days | 31.3 | 30.9 | 30.5 |

**Table A1**: Cross-correlation analysis for density-based $p$-values and prevalences of VOCs/VOIs

| | Complete | 1st truncated | 2nd truncated |
|---|---|---|---|
| Significantly dense VOCs/VOIs (VOCs only) across all countries | 61% (90%) | 64% (93%) | 63% (90%) |
| Median frequency at calling | $6 \cdot 10^{-4}$ | $4 \cdot 10^{-4}$ | $4 \cdot 10^{-4}$ |
| Median prevalence at calling | $1 \cdot 10^{-3}$ | $8 \cdot 10^{-4}$ | $1 \cdot 10^{-3}$ |
| $FD^{\text{prev}} > 0$ among variants called as significantly dense | 47% | 57% | 56% |
| $FD^{\text{des}} > 0$ among variants called as significantly dense | 47% | 52% | 49% |
| Median $FD^{\text{prev}}$ for early calls, days | 68 | 60 | 60 |
| Median $FD^{\text{des}}$ for early calls, days | 66 | 48 | 35 |
| Linear correlation and $p$-value (two-sided Student's t-test) for numbers of sequences and significantly dense VOCs/VOIs per country | 0.56 (0.024) | 0.59 (0.017) | 0.63 (0.009) |

**Table A2**: VOC/VOI calling as significantly dense subgraphs.

| | Complete | 1st truncated | 2nd truncated |
|---|---|---|---|
| VOCs/VOIs identified in at least one country | 5/10 | 5/10 | 5/10 |
| Number of countries where VOCs (VOIs) were detected | 1-15 (0) | 1-16 (0) | 1-16 (0) |
| Aggregated recall for VOCs/VOIs (VOCs only). | 19% (38%) | 22% (44%) | 21% (41%) |
| Precision: percentage of densest communities with at least 80% similarity with VOCs/VOIs, aggregated across all countries. | 31% | 21% | 16% |
| Percentage of earliest VOCs/VOIs detections with $FD^{\text{prev}} \geq 0$ | 40% | 43% | 30% |
| Percentage of earliest VOCs/VOIs detections with $FD^{\text{des}} \geq 0$ | 67% | 40% | 33% |
| Median cumulative frequency at first detection | $1.4 \cdot 10^{-3}$ | $9.96 \cdot 10^{-4}$ | $1.5 \cdot 10^{-3}$ |
| Median prevalence at first detection | $1.59 \cdot 10^{-2}$ | $1.75 \cdot 10^{-2}$ | $2.45 \cdot 10^{-2}$ |
| VOCs/VOIs (VOCs) with $FD^{\text{prev}} \geq 0$ in at least one country | 5/10 (5/5) | 5/10 (5/5) | 5/10 (5/5) |
| VOCs/VOIs (VOCs) with $FD^{\text{des}} \geq 0$ in at least one country | 5/10 (5/5) | 5/10 (5/5) | 5/10 (5/5) |
| Median $FD^{\text{prev}}$ for early calls, days | 218 | 30 | 30 |
| Median $FD^{\text{des}}$ for early calls, days | 123 | 44 | 36 |

**Table A3**: Analysis of densest subnetworks

#### A.2 Analysis of densest subnetworks of coordinated substitution networks

The results are summarized in Table A3 and Figs. 5, A33-A40. The main findings here can be summarized as follows:

- Percentages of densest communities that were at least 80% identical to the known variants ranged from 16% in the second truncated dataset to 31% in the complete dataset (Table A3). All detected variants were VOCs.

- Among these communities, 33% − 67% were early detected before the corresponding VOCs received official designation from WHO. Similarly, 30% − 40% were early detected before the VOCs attained a 1% prevalence (see Figs. A38-A40 and Table A3).
- For the three datasets, the median cumulative frequencies of VOCs at times of their early detection ranged from $5 \cdot 10^{-4}$ to $9 \cdot 10^{-4}$. Respective median prevalences were between $3 \cdot 10^{-3}$ and $9 \cdot 10^{-3}$ (refer to Figs. A38-A40 and Table A3).
- Across all datasets, every VOC was detected with a minimum 0.8 accuracy in at least one country before its official designation (see Figs. A38, A36-A37).
- Maximal forecasting depths $FD^{\text{des}}$ relative to the WHO designation were $231, 111, 150, 270, 319$ days (complete dataset), $231, 111, 135, 285, 319$ days (first truncated dataset), $36, 111, 45, 285, 4$ days (second truncated dataset)(Figs. A38-A40). Median depths varied between $FD^{\text{des}} = 123$ for the complete dataset and $FD^{\text{des}} = 44$ and $FD^{\text{des}} = 36$ for truncated datasets.
- The numbers for early detection in relation to the 1% prevalence were similar, except for the Beta variant. This was detected as soon as it reached the 1% mark ($FD^{\text{prev}} = 0$) (see Figs. A38-A40).

#### A.3 Supplementary figures

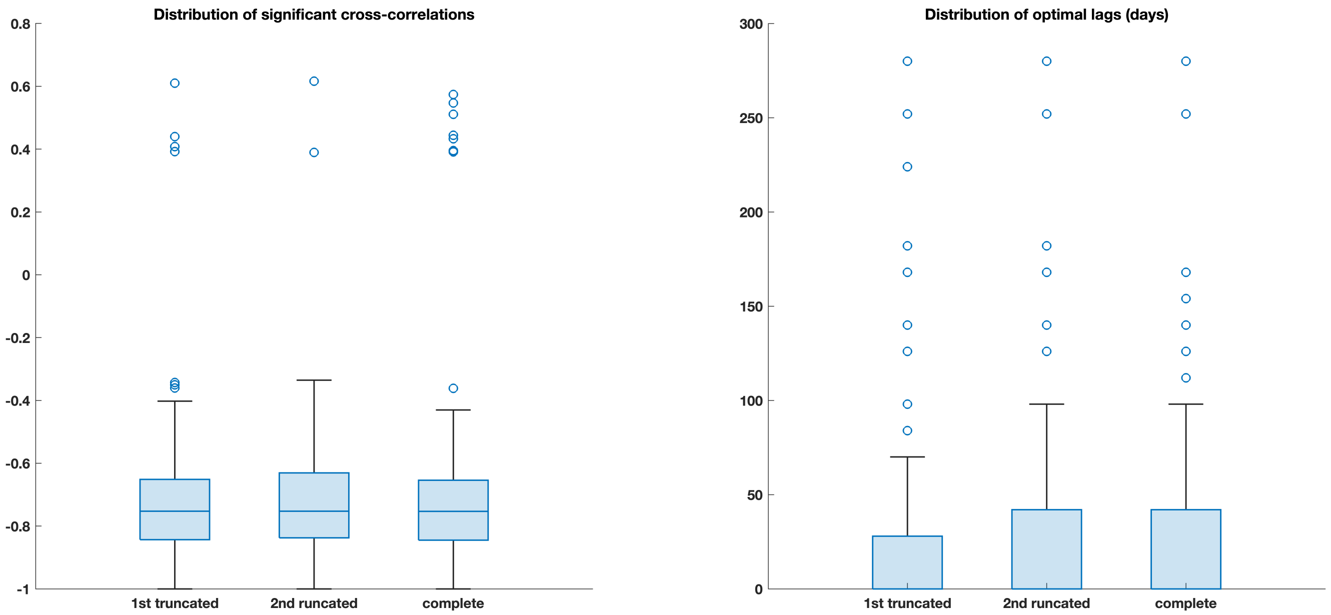


**Fig. A1**: Distributions of statistically significant cross-correlations and optimal lags for three analyzed datasets

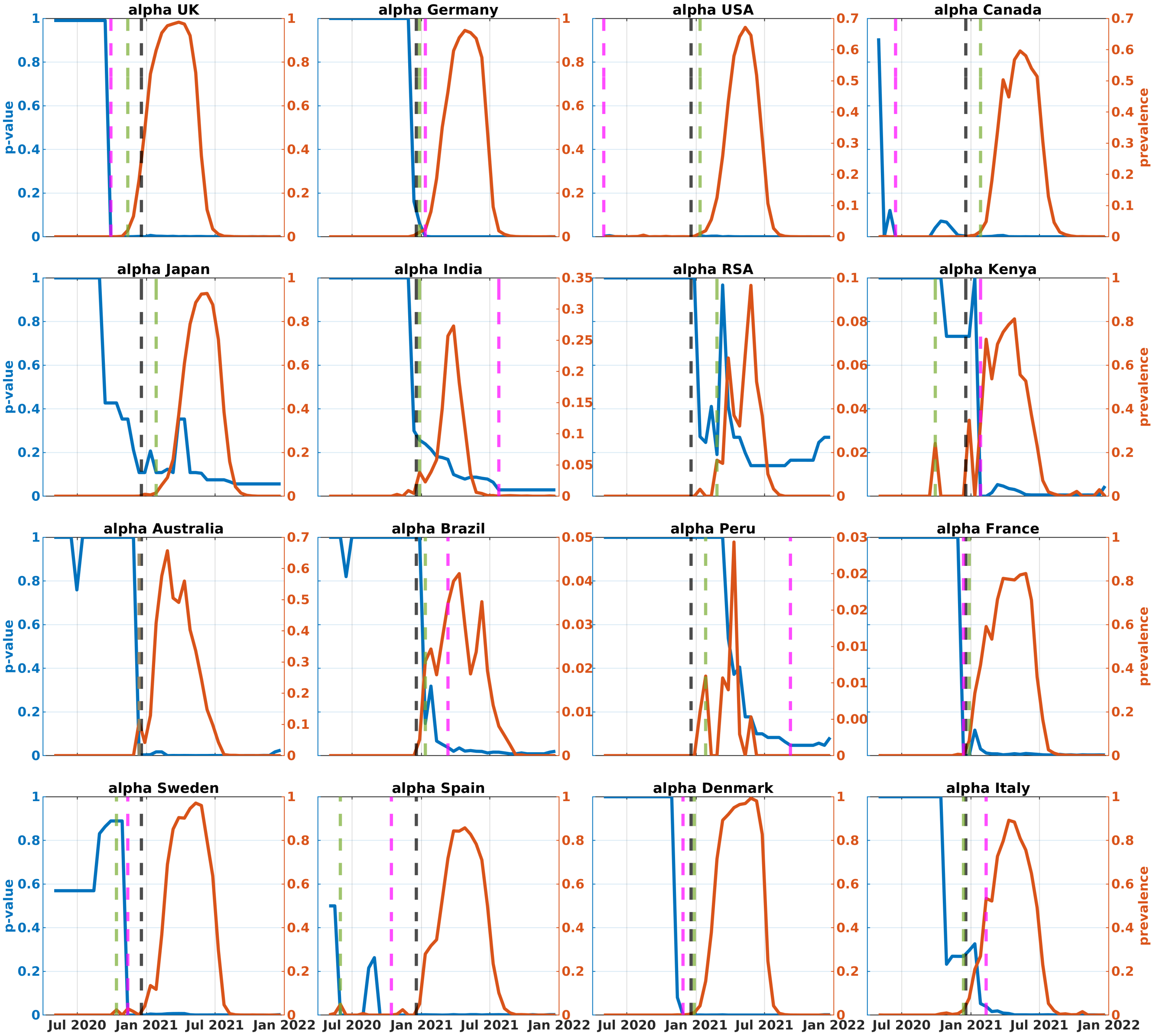


**Fig. A2**: $p$-values (blue) and prevalences (red) of Alpha variant in the analyzed countries (complete dataset). Black, green, and magenta lines represent the times of VOC designation, achieving 1% prevalence, and becoming significantly dense, respectively.

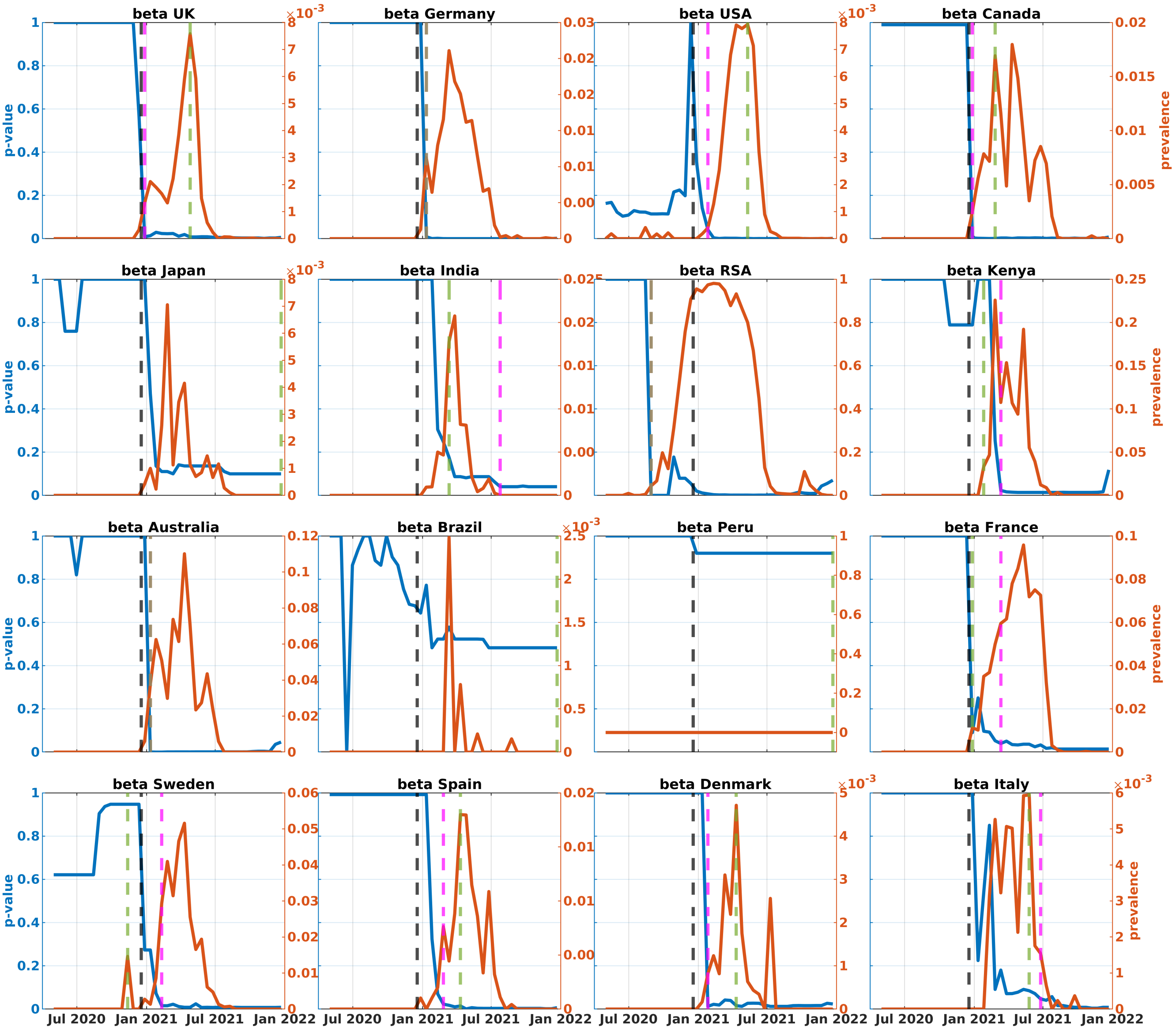


**Fig. A3**: $p$-values (blue) and prevalences (red) of Beta variant in the analyzed countries (complete dataset). Black, green, and magenta lines represent the times of VOC designation, achieving 1% prevalence, and becoming significantly dense, respectively.

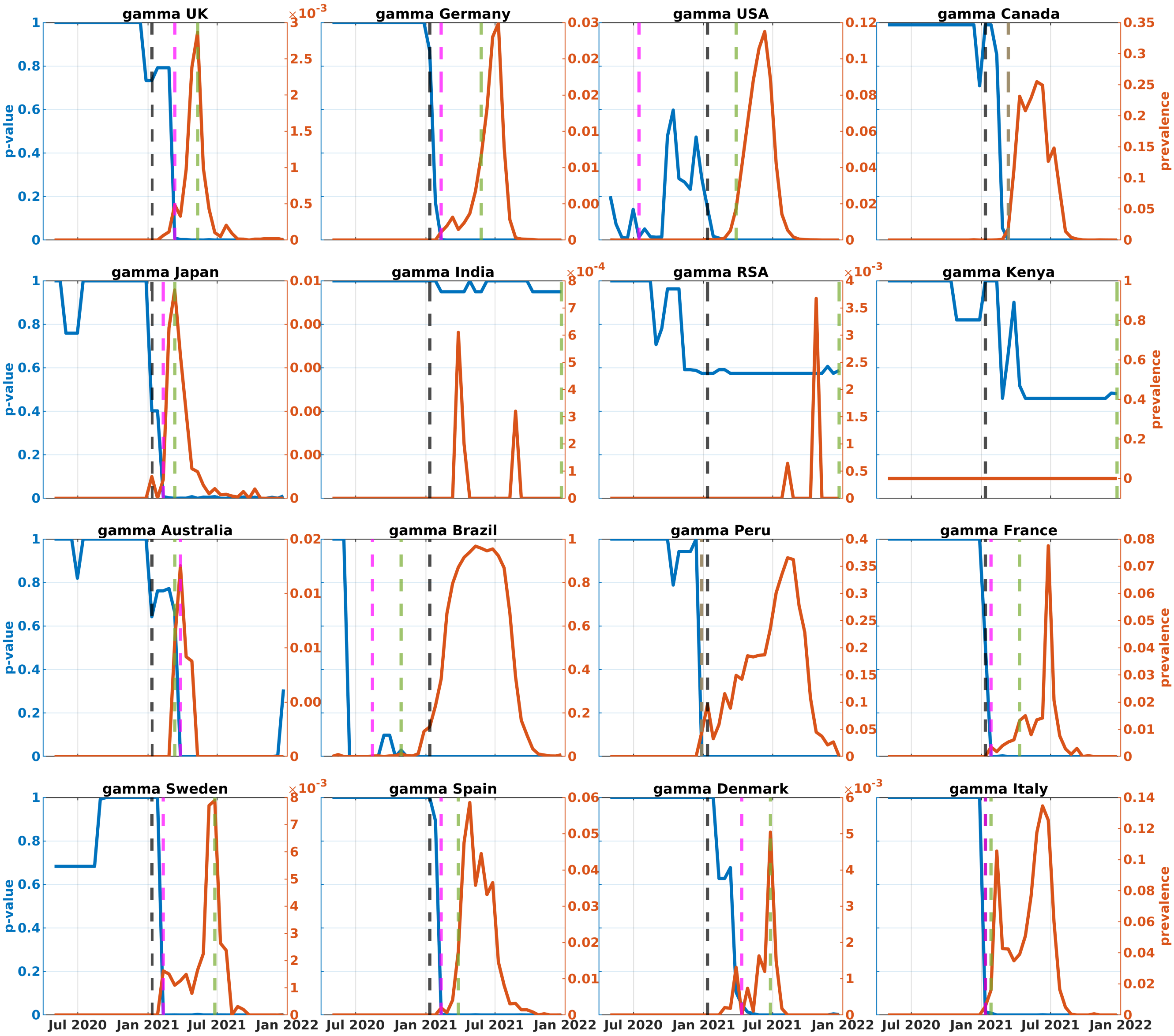


**Fig. A4**: $p$-values (blue) and prevalences (red) of Gamma variant in the analyzed countries (complete dataset). Black, green, and magenta lines represent the times of VOC designation, achieving 1% prevalence, and becoming significantly dense, respectively.

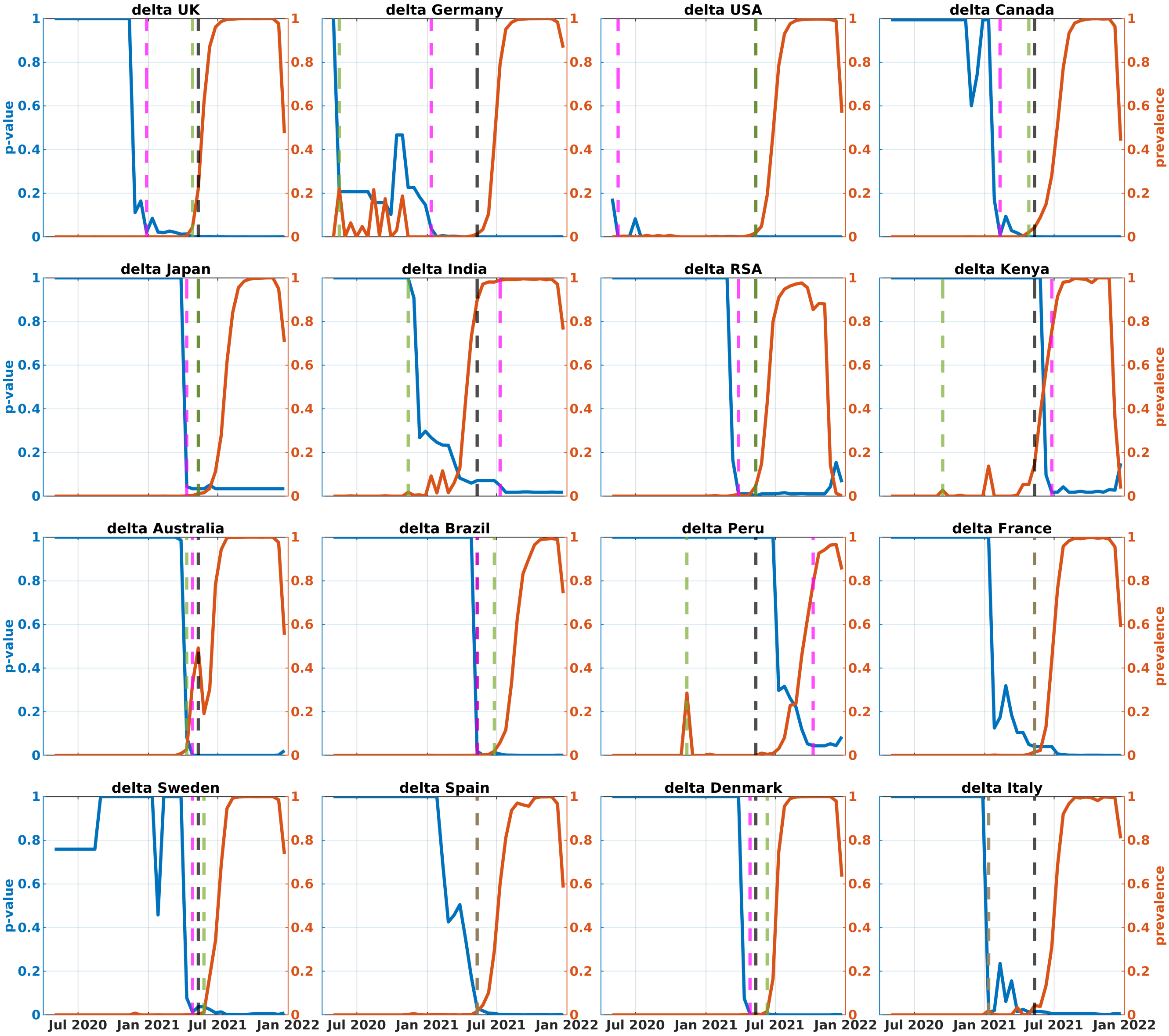


**Fig. A5**: $p$-values (blue) and prevalences (red) of Delta variant in the analyzed countries (complete dataset). Black, green, and magenta lines represent the times of VOC designation, achieving 1% prevalence, and becoming significantly dense, respectively.

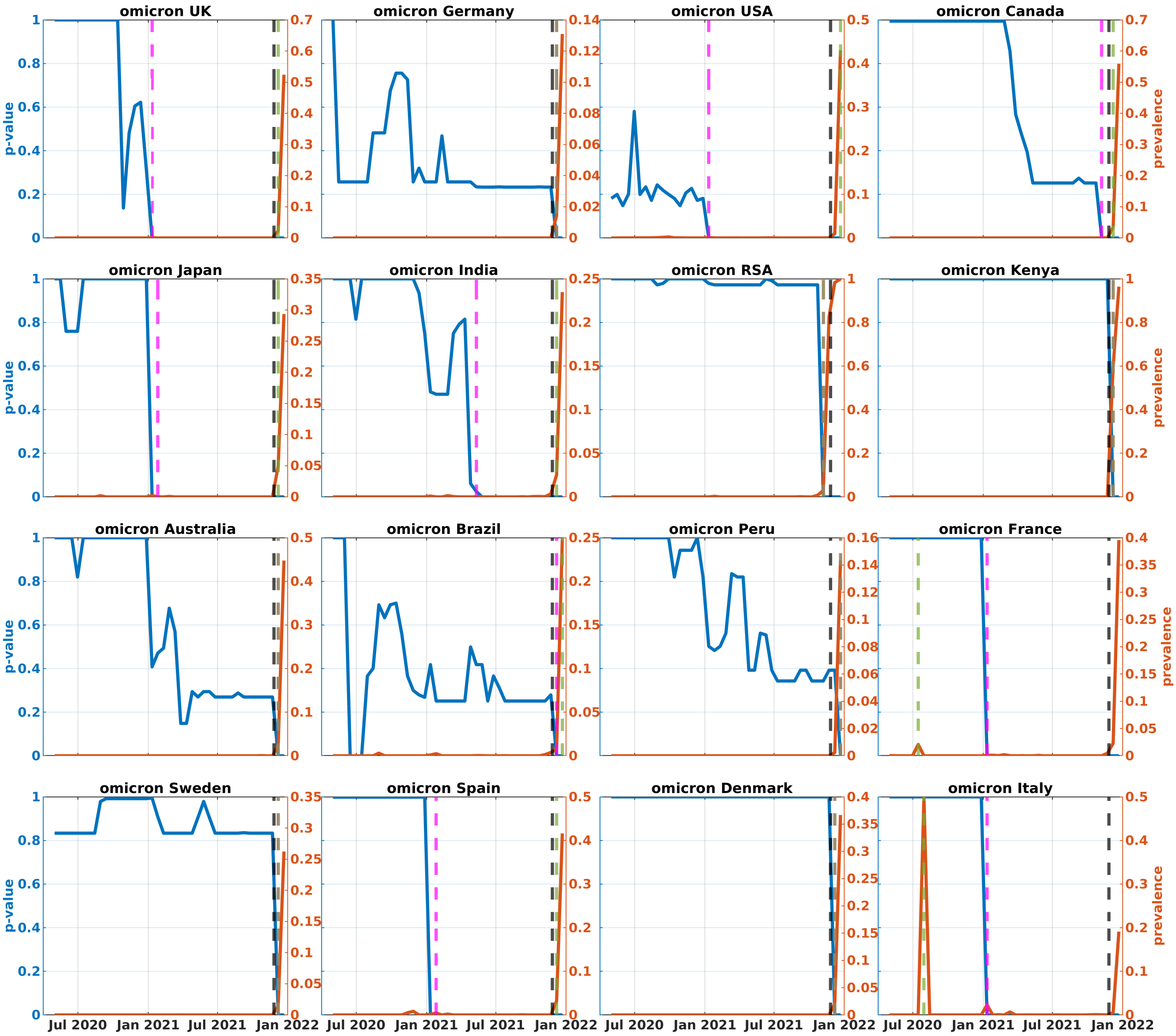


**Fig. A6**: $p$-values (blue) and prevalences (red) of Omicron variant in the analyzed countries (complete dataset). Black, green, and magenta lines represent the times of VOC designation, achieving 1% prevalence, and becoming significantly dense, respectively.

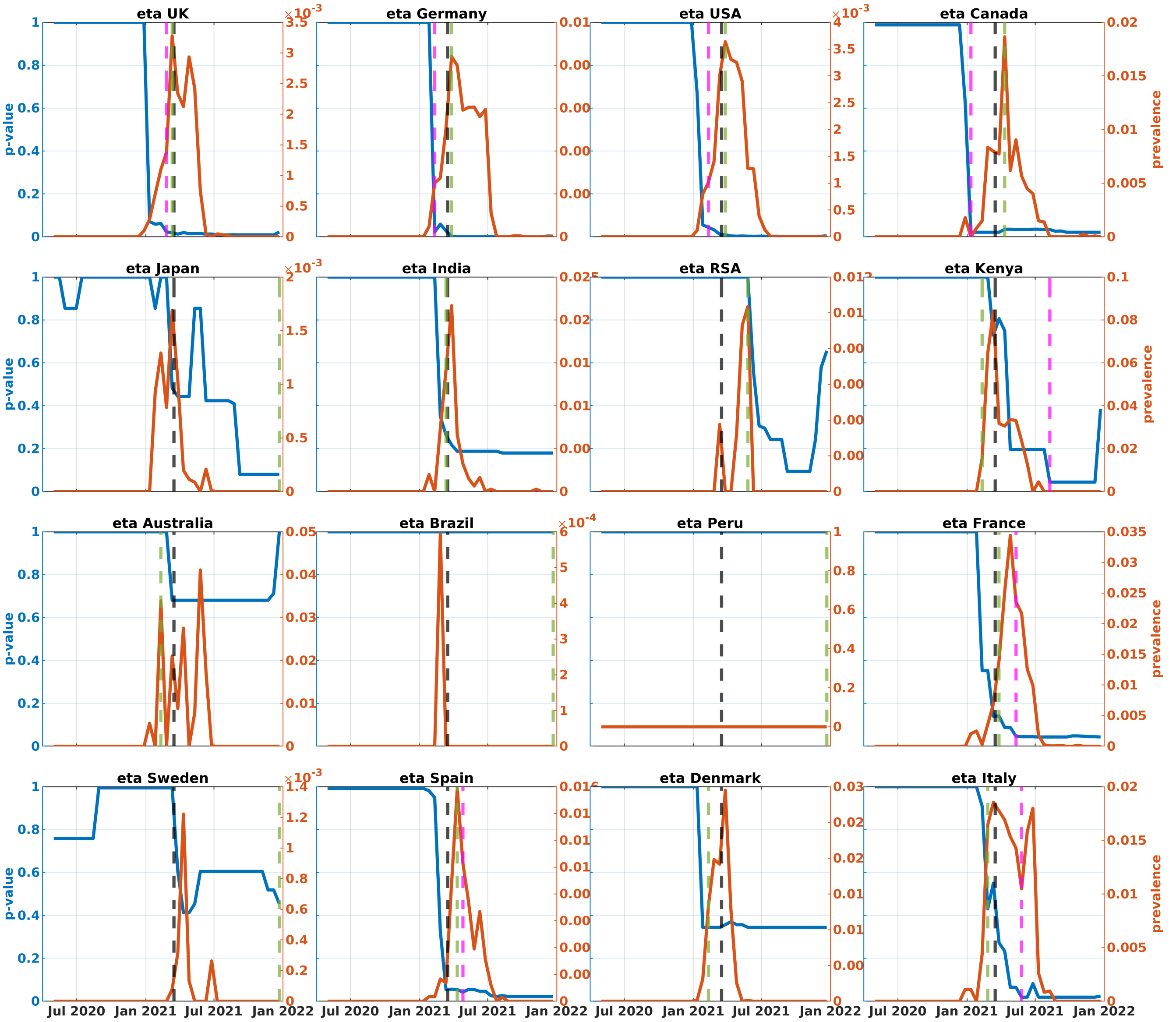


**Fig. A7**: $p$-values (blue) and prevalences (red) of Eta variant in the analyzed countries (complete dataset). Black, green, and magenta lines represent the times of VOC designation, achieving 1% prevalence, and becoming significantly dense, respectively.

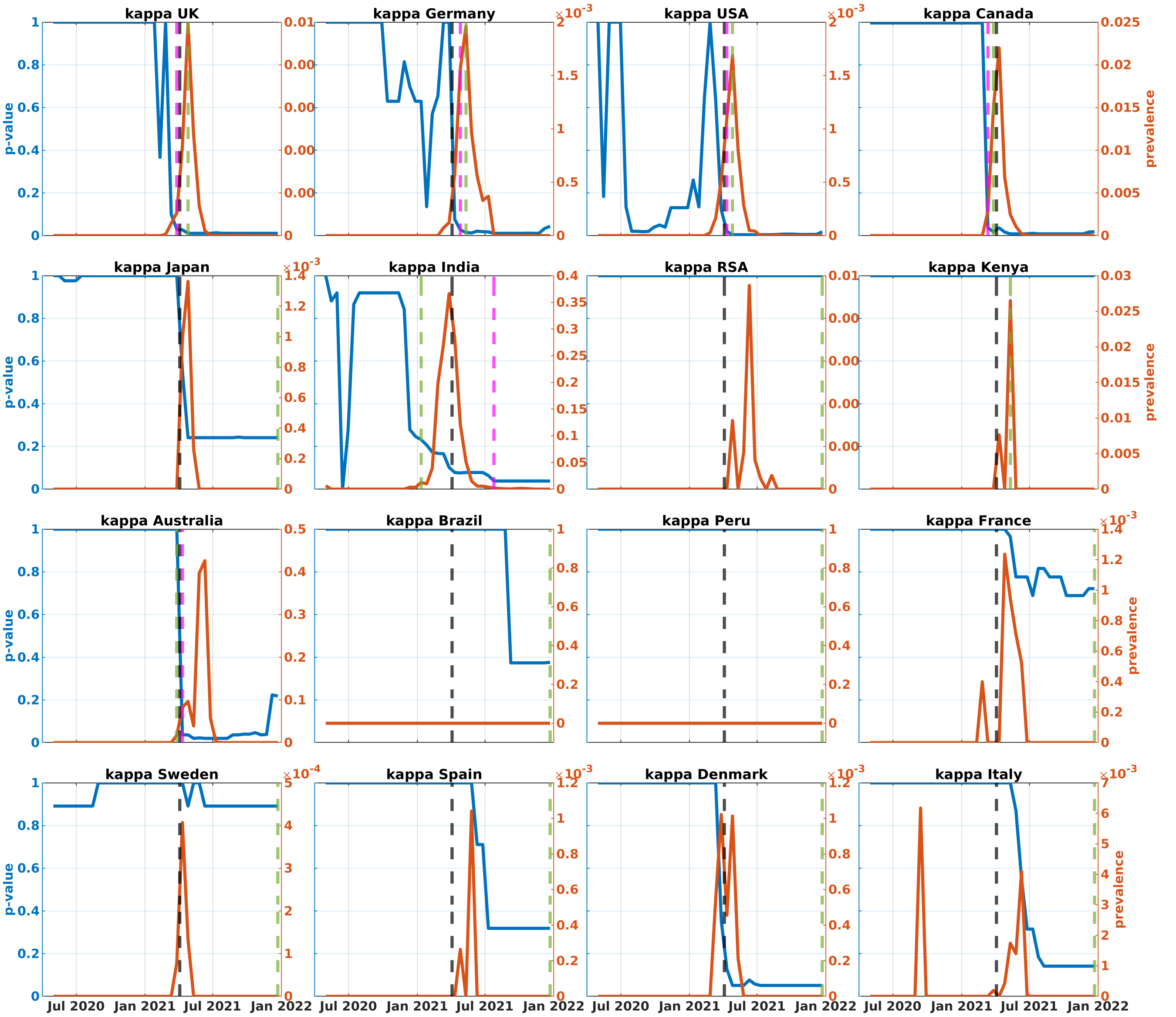


**Fig. A8**: $p$-values (blue) and prevalences (red) of Kappa variant in the analyzed countries (complete dataset). Black, green, and magenta lines represent the times of VOC designation, achieving 1% prevalence, and becoming significantly dense, respectively.

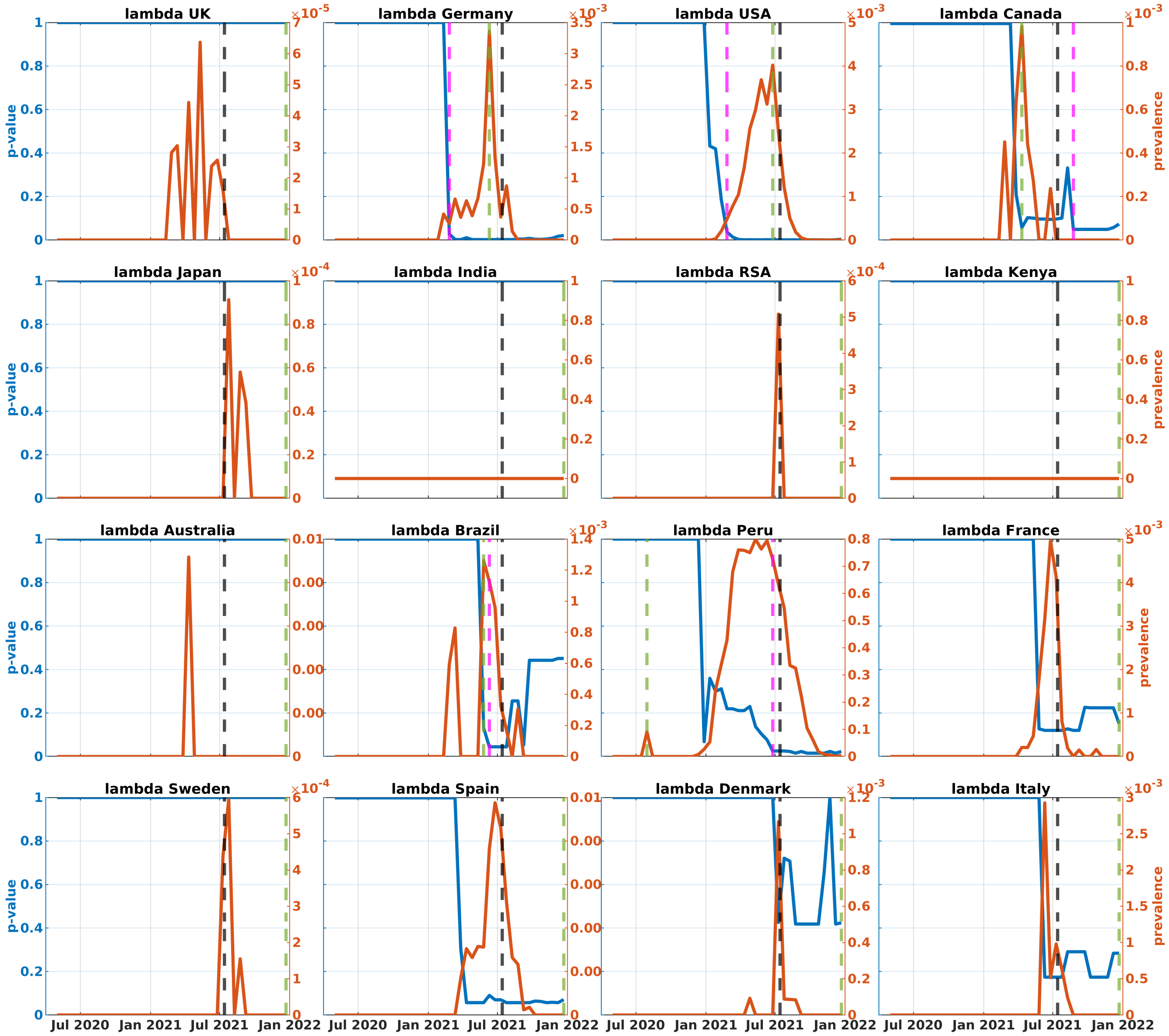


**Fig. A9**: $p$-values (blue) and prevalences (red) of Lambda variant in the analyzed countries (complete dataset). Black, green, and magenta lines represent the times of VOC designation, achieving 1% prevalence, and becoming significantly dense, respectively.

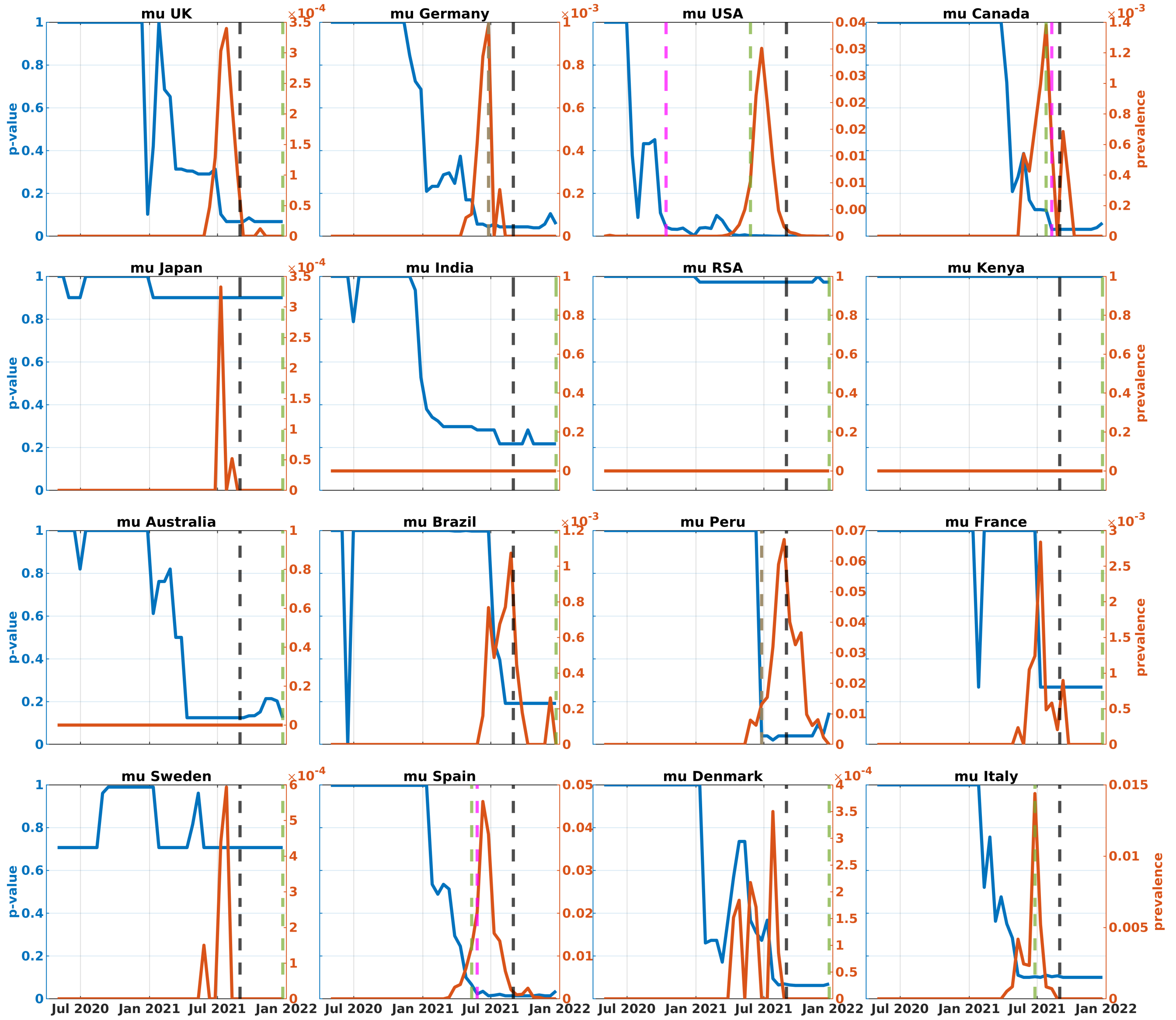


**Fig. A10**: $p$-values (blue) and prevalences (red) of Mu variant in the analyzed countries (complete dataset). Black, green, and magenta lines represent the times of VOC designation, achieving 1% prevalence, and becoming significantly dense, respectively.

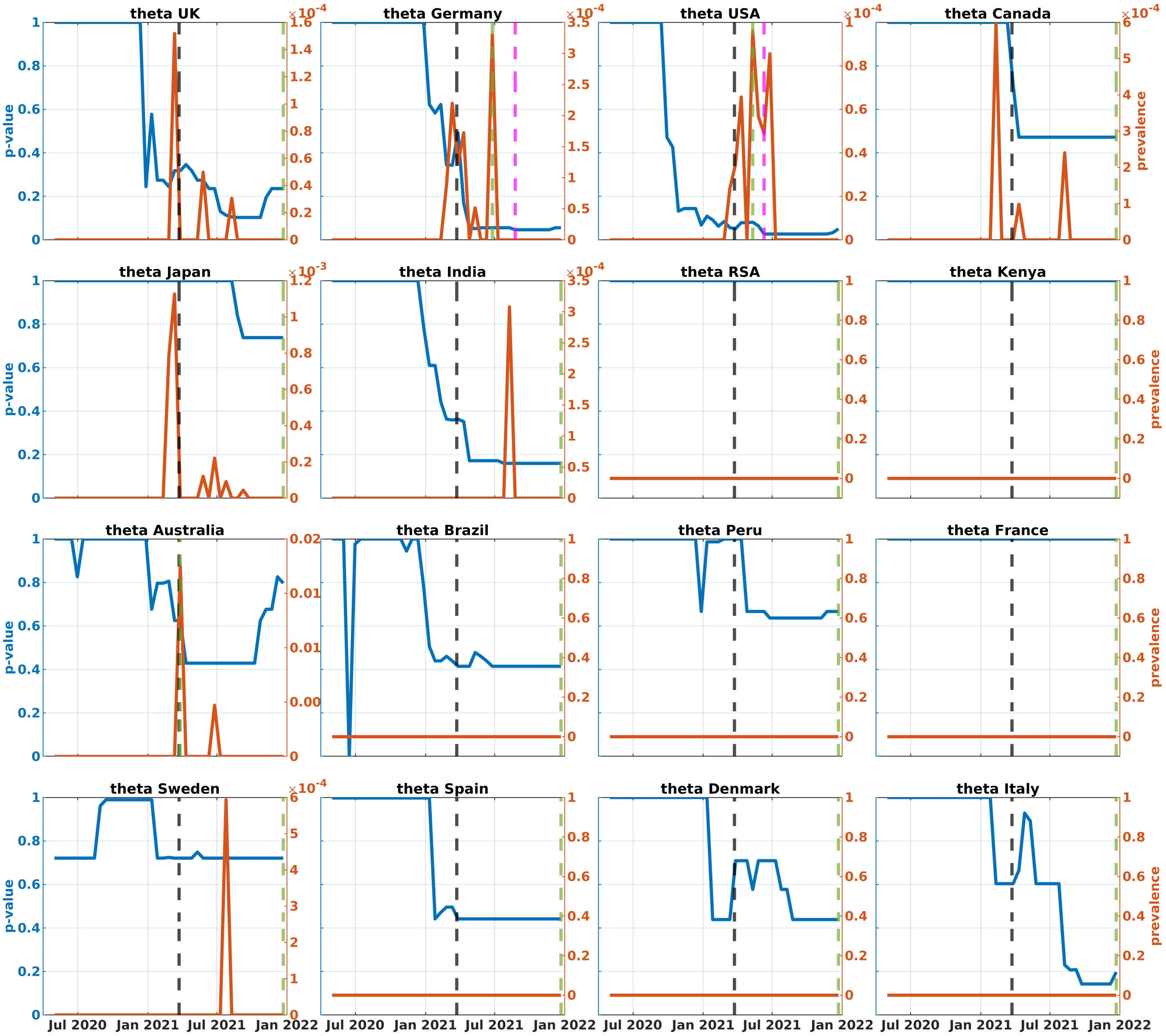


**Fig. A11**: $p$-values (blue) and prevalences (red) of Theta variant in the analyzed countries (complete dataset). Black, green, and magenta lines represent the times of VOC designation, achieving 1% prevalence, and becoming significantly dense, respectively.

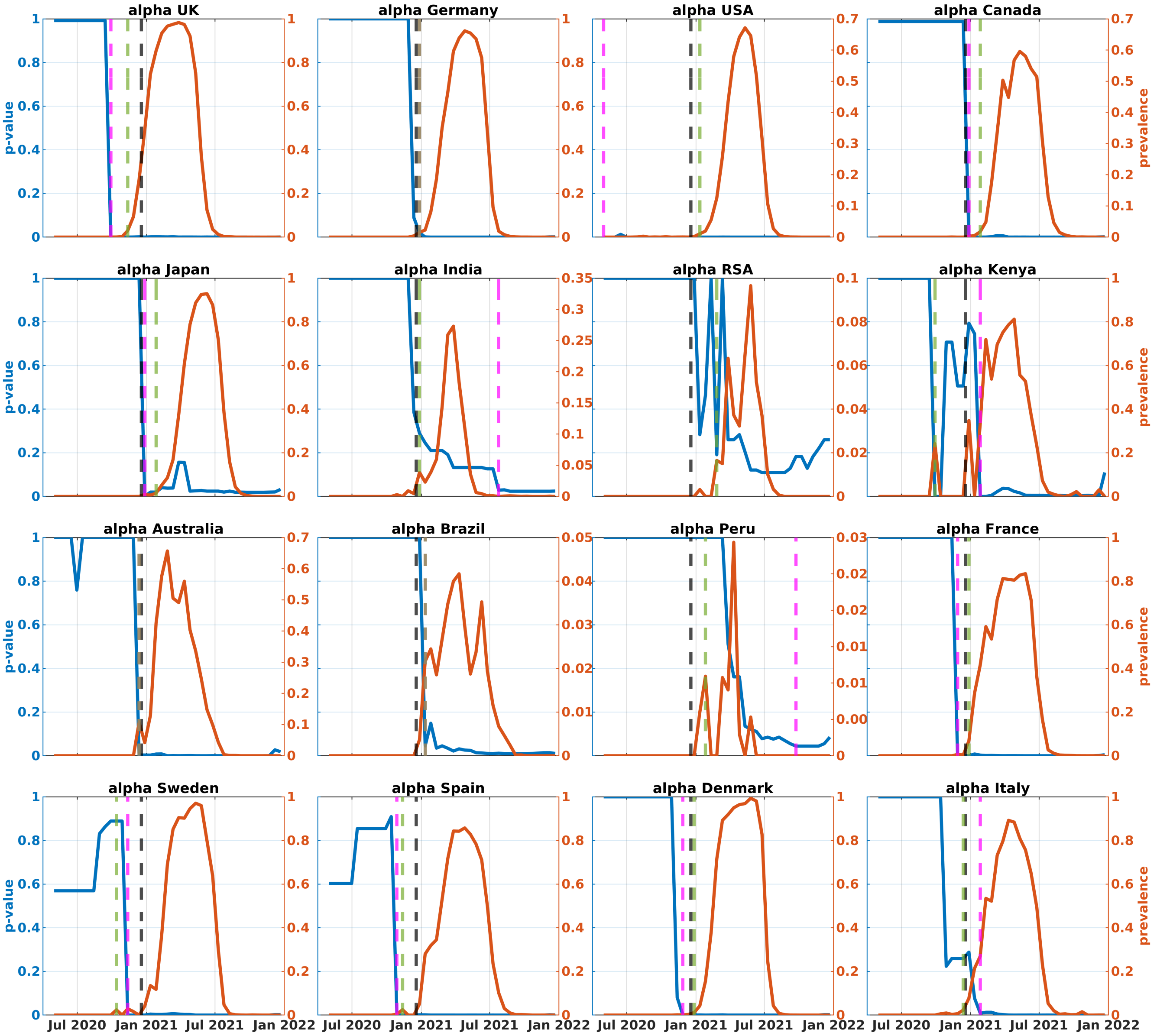


**Fig. A12**: $p$-values (blue) and prevalences (red) of Alpha variant in the analyzed countries (first truncated dataset). Black, green, and magenta lines represent the times of VOC designation, achieving 1% prevalence, and becoming significantly dense, respectively.

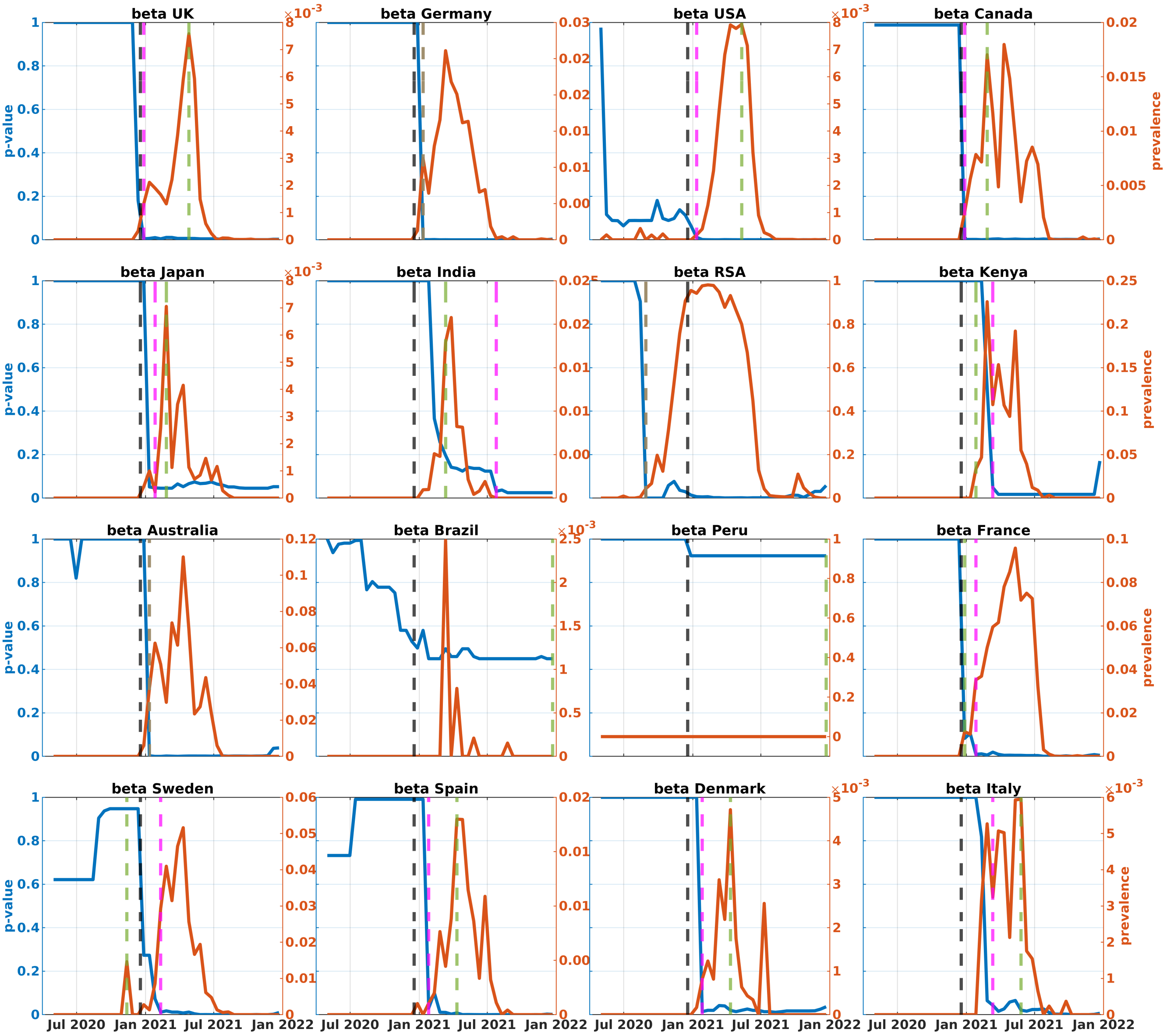

**Fig. A13**: $p$-values (blue) and prevalences (red) of Beta variant in the analyzed countries (first truncated dataset). Black, green, and magenta lines represent the times of VOC designation, achieving 1% prevalence, and becoming significantly dense, respectively.

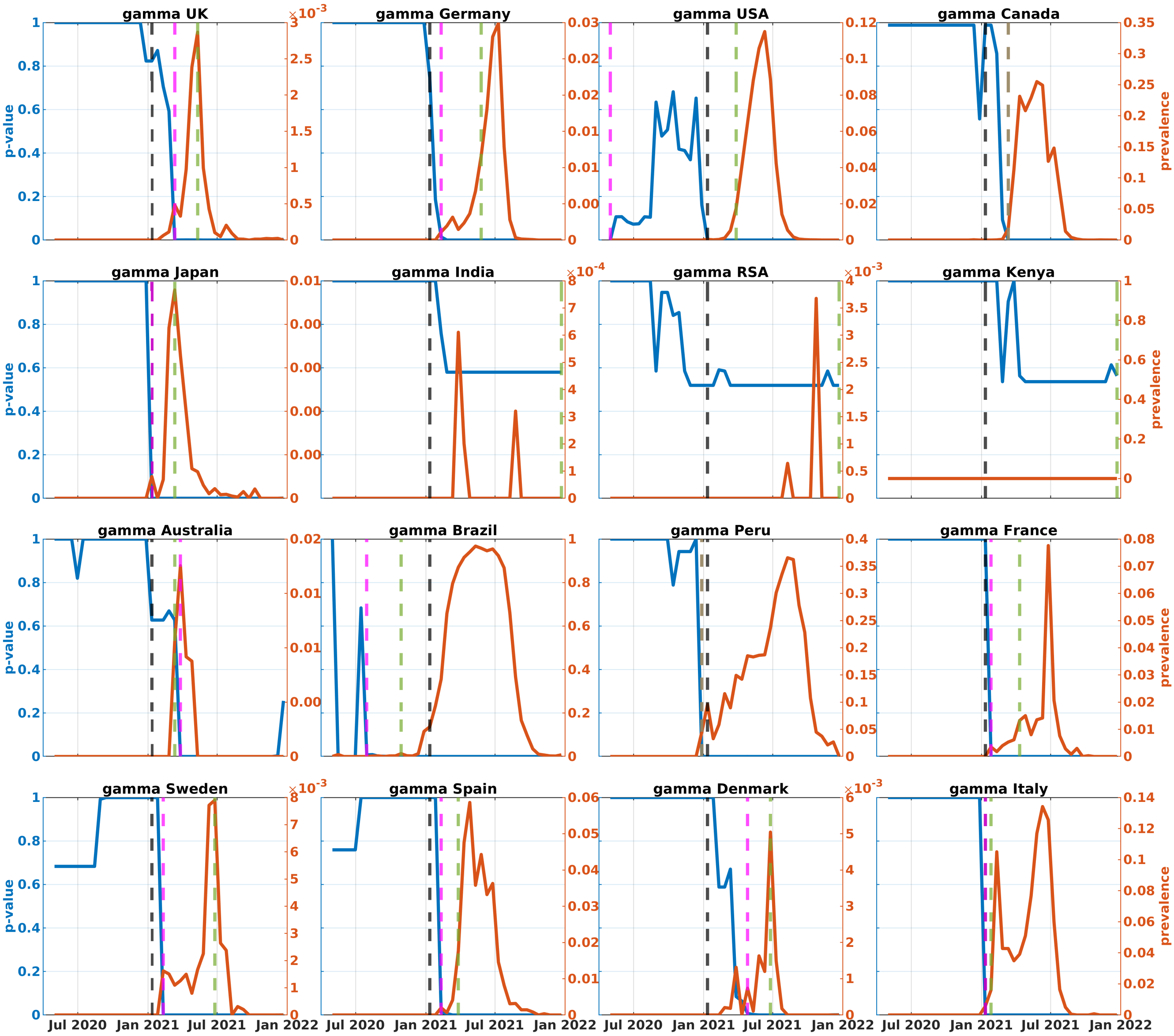


**Fig. A14**: $p$-values (blue) and prevalences (red) of Gamma variant in the analyzed countries (first truncated dataset). Black, green, and magenta lines represent the times of VOC designation, achieving 1% prevalence, and becoming significantly dense, respectively.

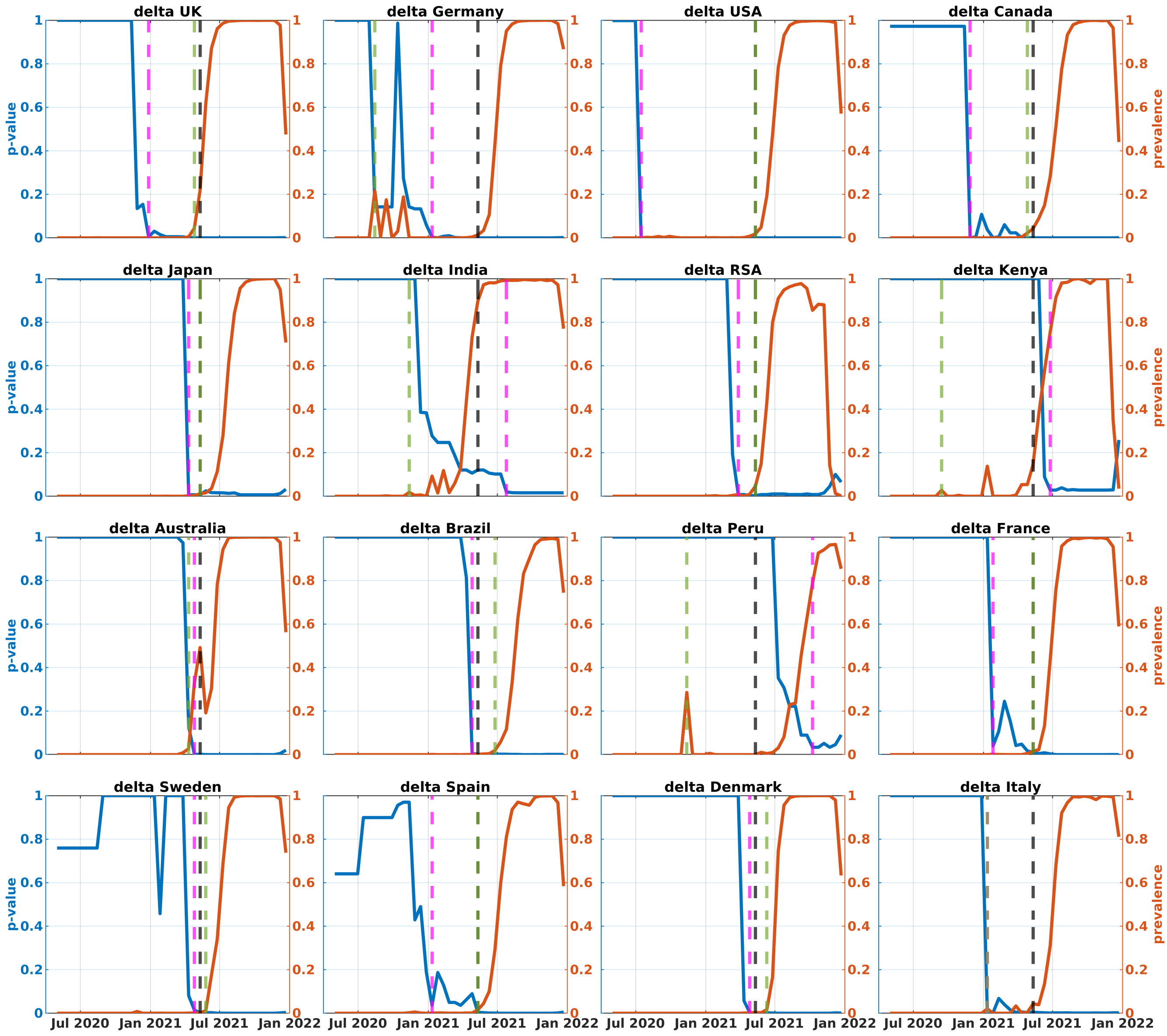


**Fig. A15**: $p$-values (blue) and prevalences (red) of Delta variant in the analyzed countries (first truncated dataset). Black, green, and magenta lines represent the times of VOC designation, achieving 1% prevalence, and becoming significantly dense, respectively.

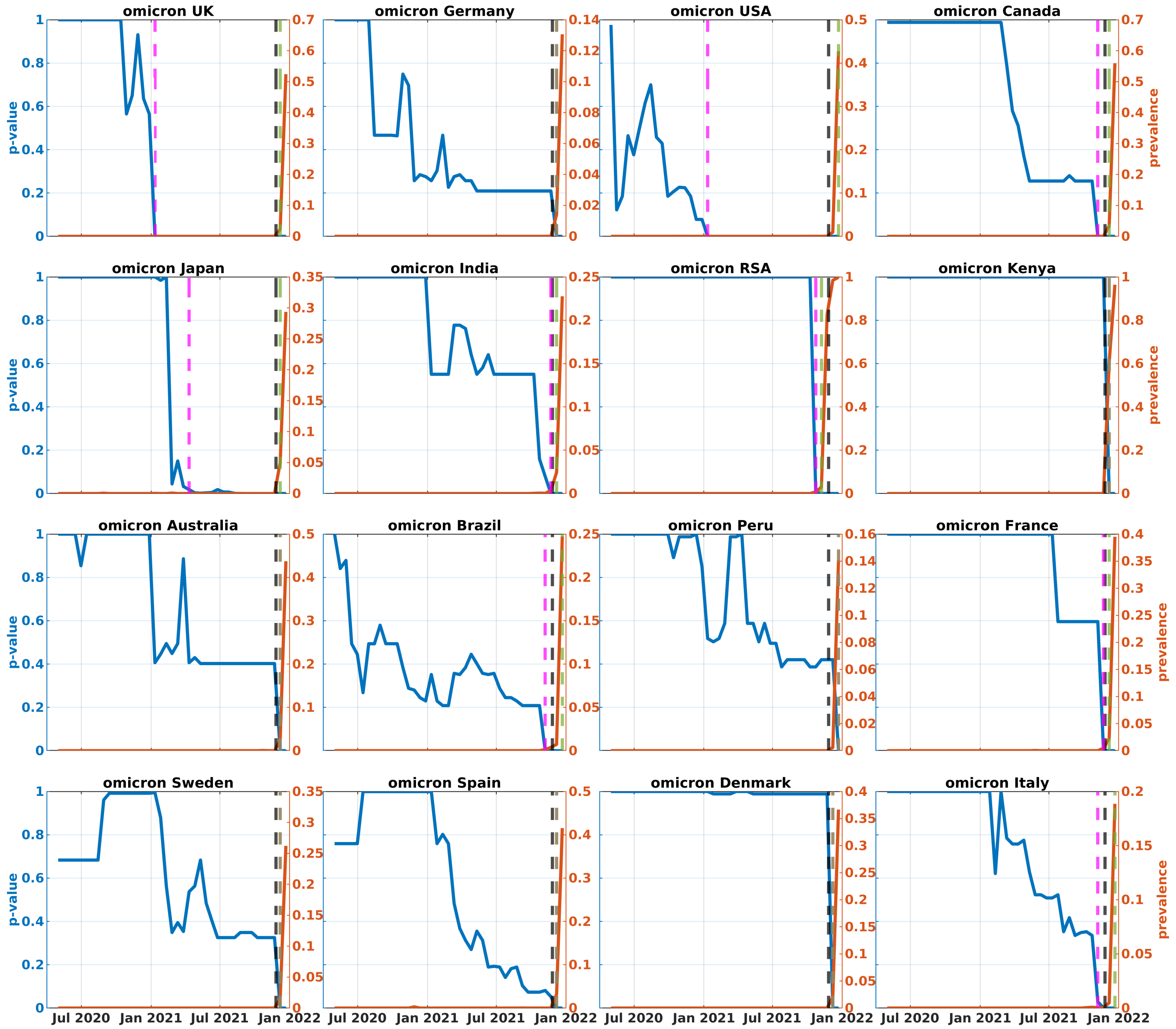


**Fig. A16**: $p$-values (blue) and prevalences (red) of Omicron variant in the analyzed countries (first truncated dataset). Black, green, and magenta lines represent the times of VOC designation, achieving 1% prevalence, and becoming significantly dense, respectively.

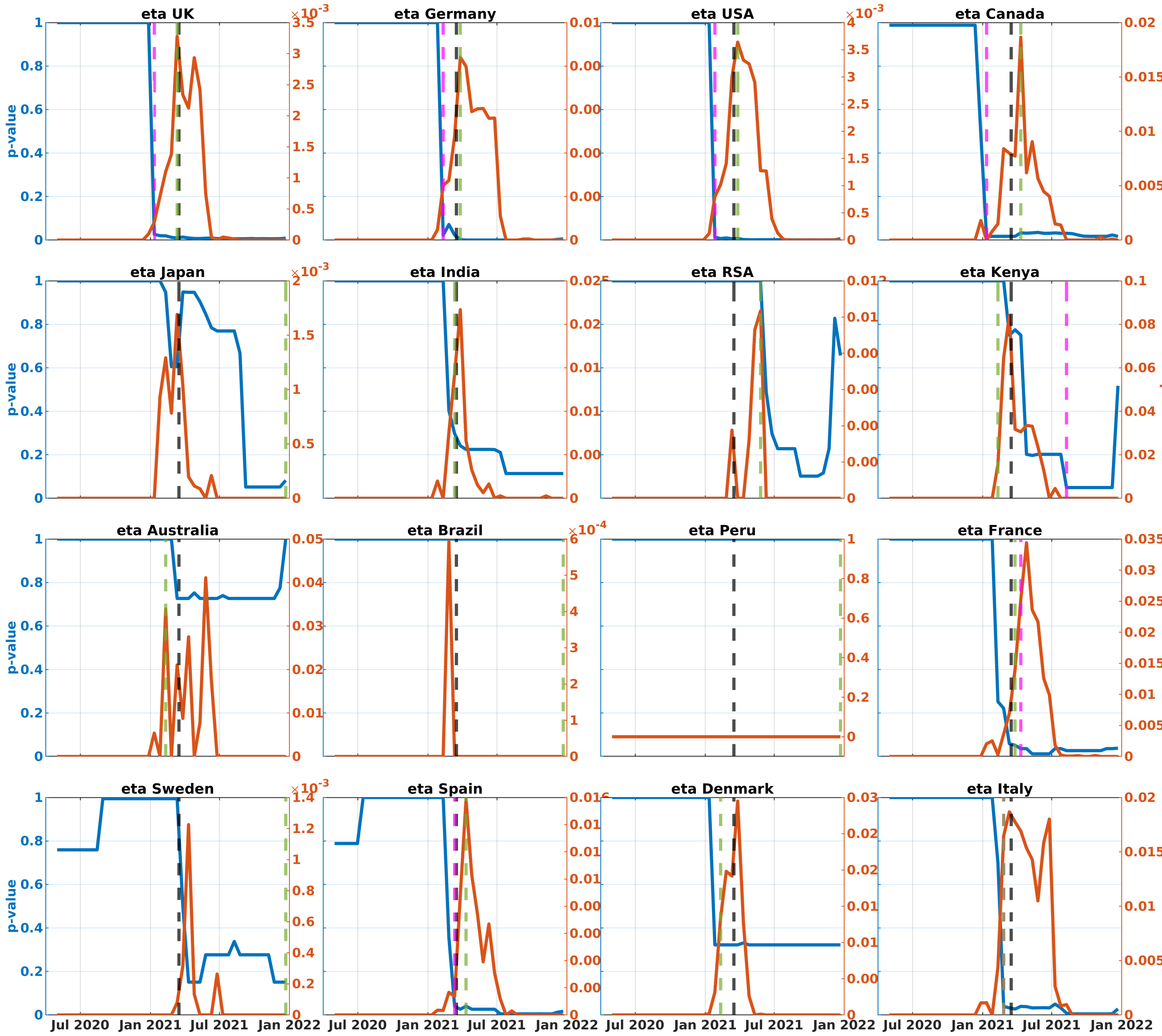


**Fig. A17**: $p$-values (blue) and prevalences (red) of Eta variant in the analyzed countries (first truncated dataset). Black, green, and magenta lines represent the times of VOC designation, achieving 1% prevalence, and becoming significantly dense, respectively.

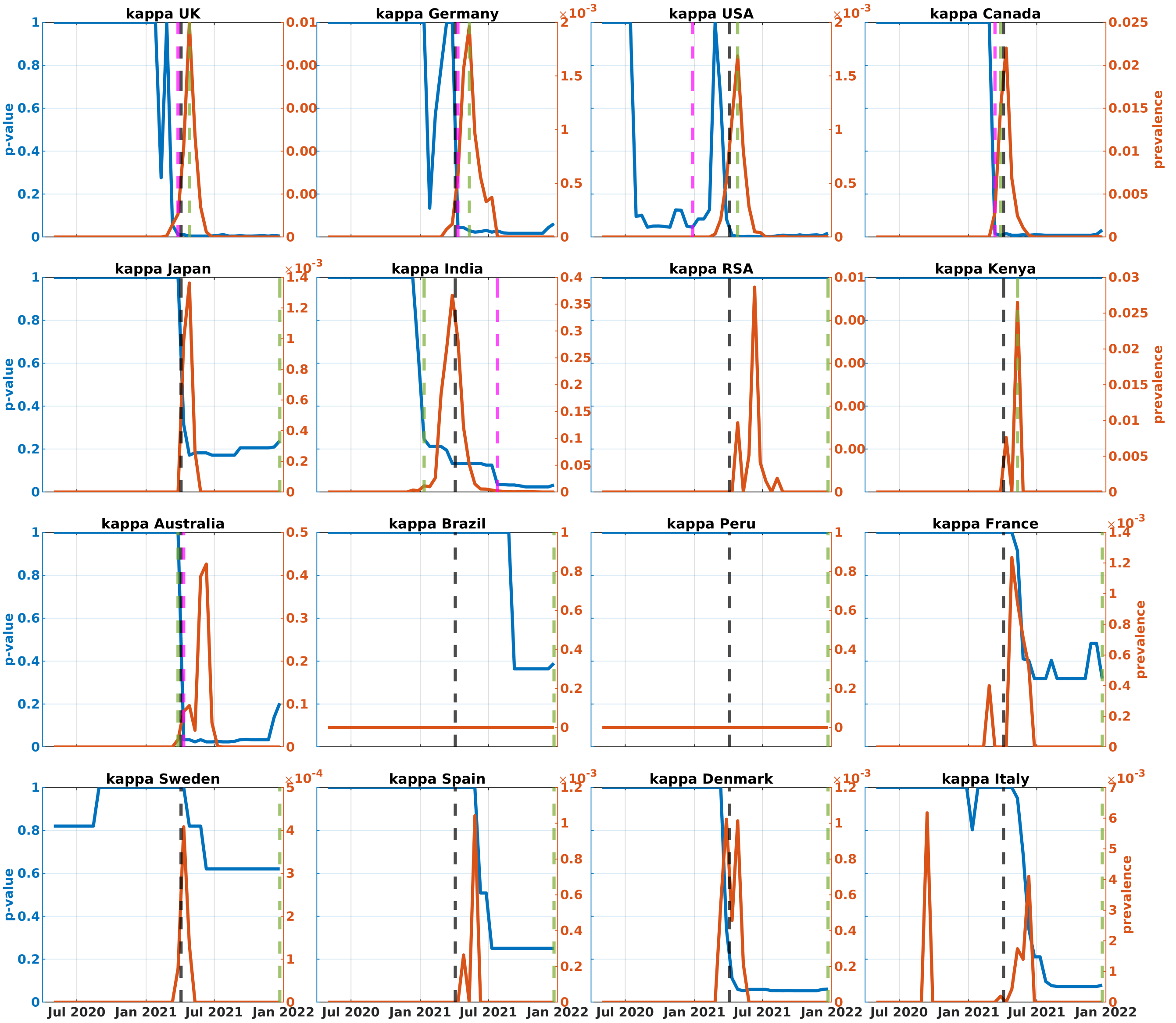


**Fig. A18**: $p$-values (blue) and prevalences (red) of Kappa variant in the analyzed countries (first truncated dataset). Black, green, and magenta lines represent the times of VOC designation, achieving 1% prevalence, and becoming significantly dense, respectively.

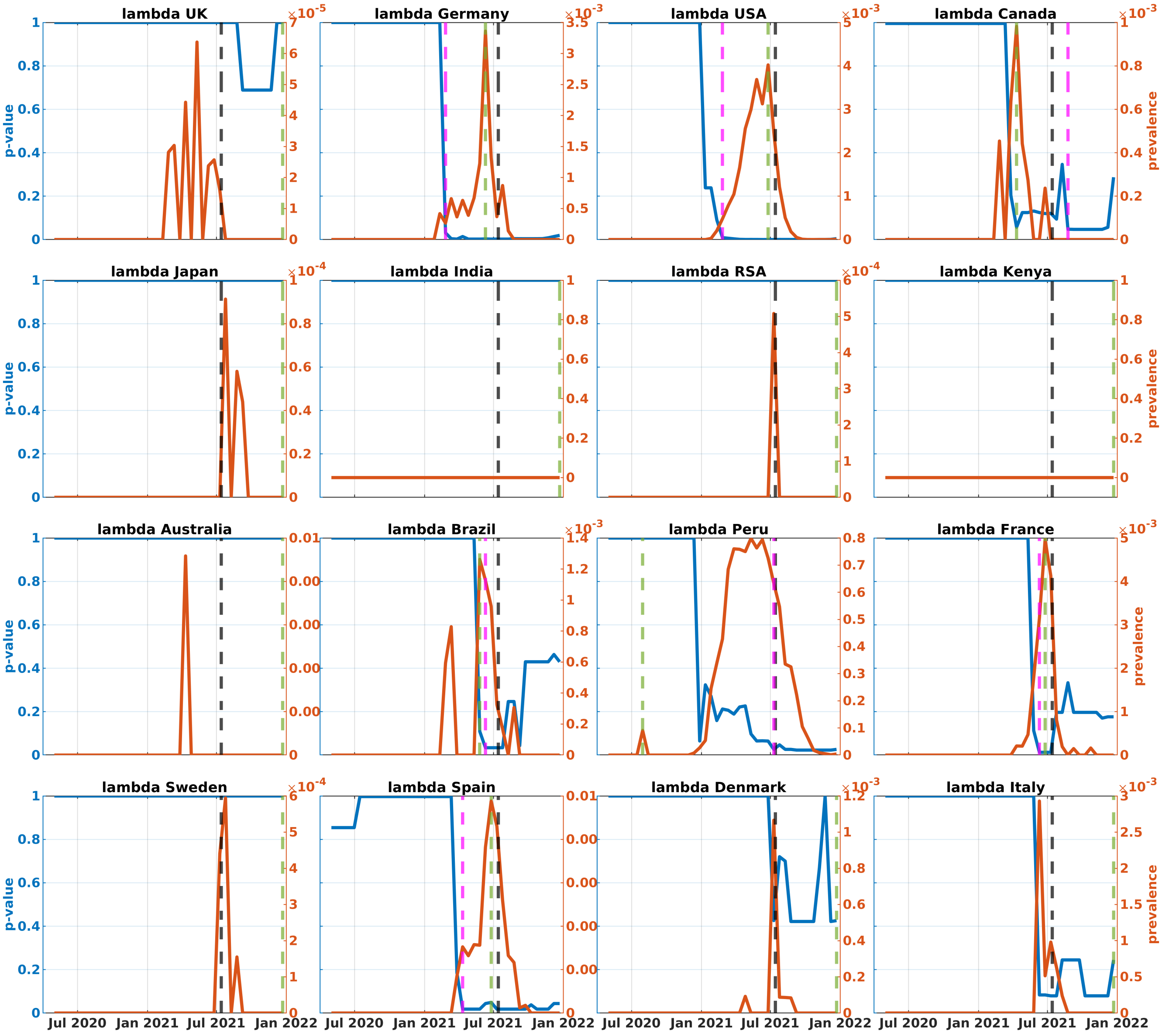


**Fig. A19**: $p$-values (blue) and prevalences (red) of Lambda variant in the analyzed countries (first truncated dataset). Black, green, and magenta lines represent the times of VOC designation, achieving 1% prevalence, and becoming significantly dense, respectively.

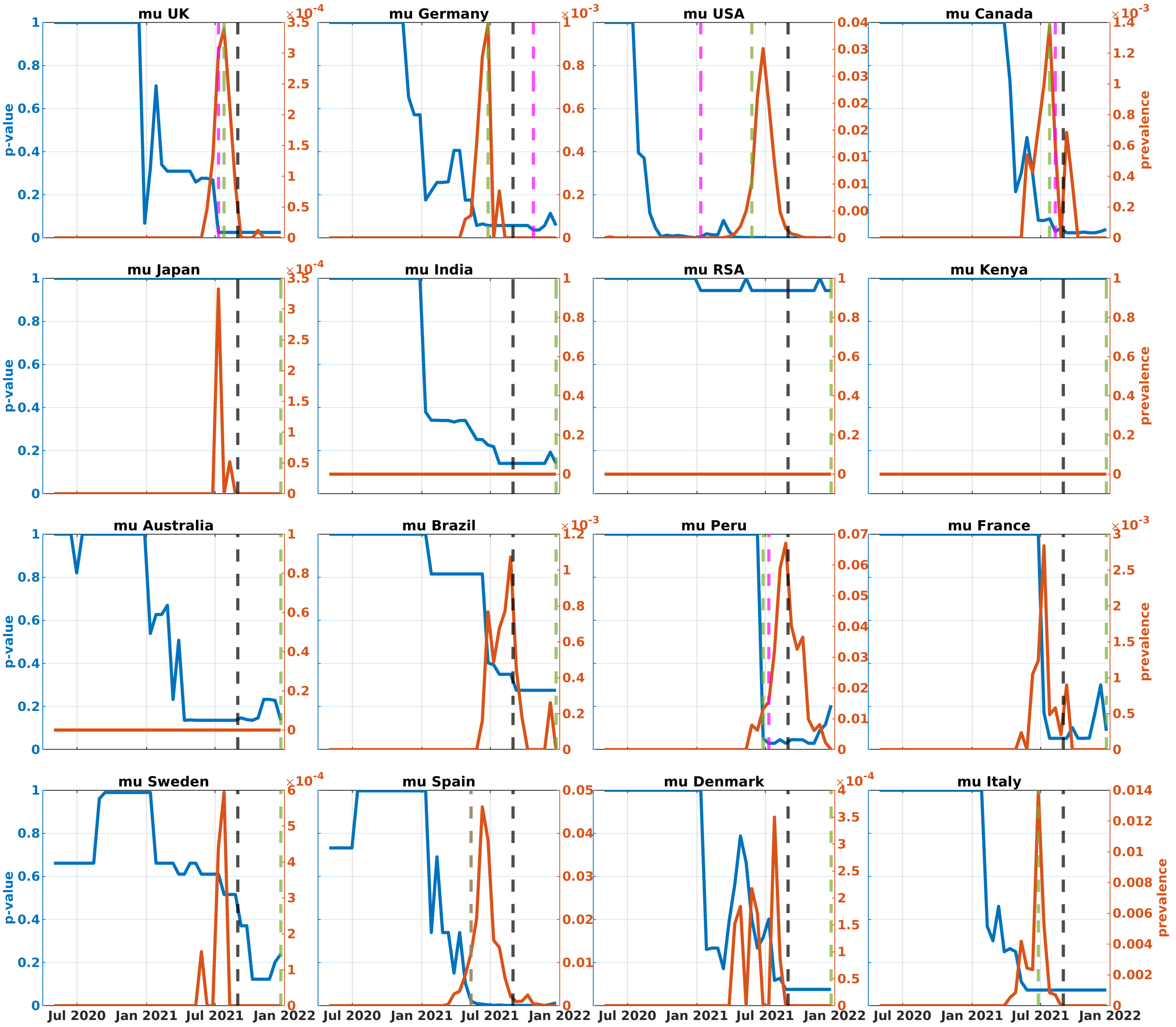


**Fig. A20**: $p$-values (blue) and prevalences (red) of Mu variant in the analyzed countries (first truncated dataset). Black, green, and magenta lines represent the times of VOC designation, achieving 1% prevalence, and becoming significantly dense, respectively.

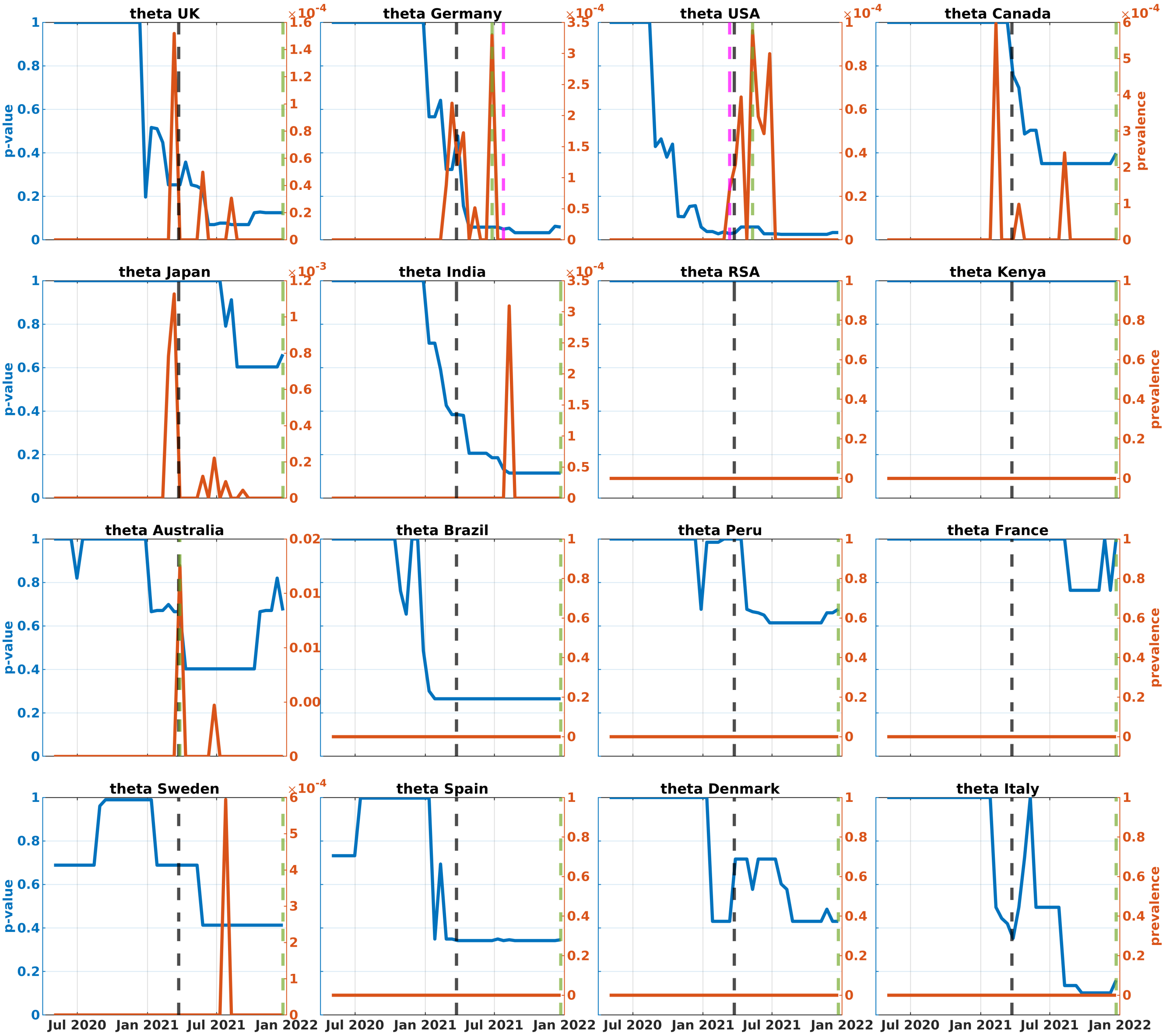

**Fig. A21**: $p$-values (blue) and prevalences (red) of Theta variant in the analyzed countries (first truncated dataset). Black, green, and magenta lines represent the times of VOC designation, achieving 1% prevalence, and becoming significantly dense, respectively.

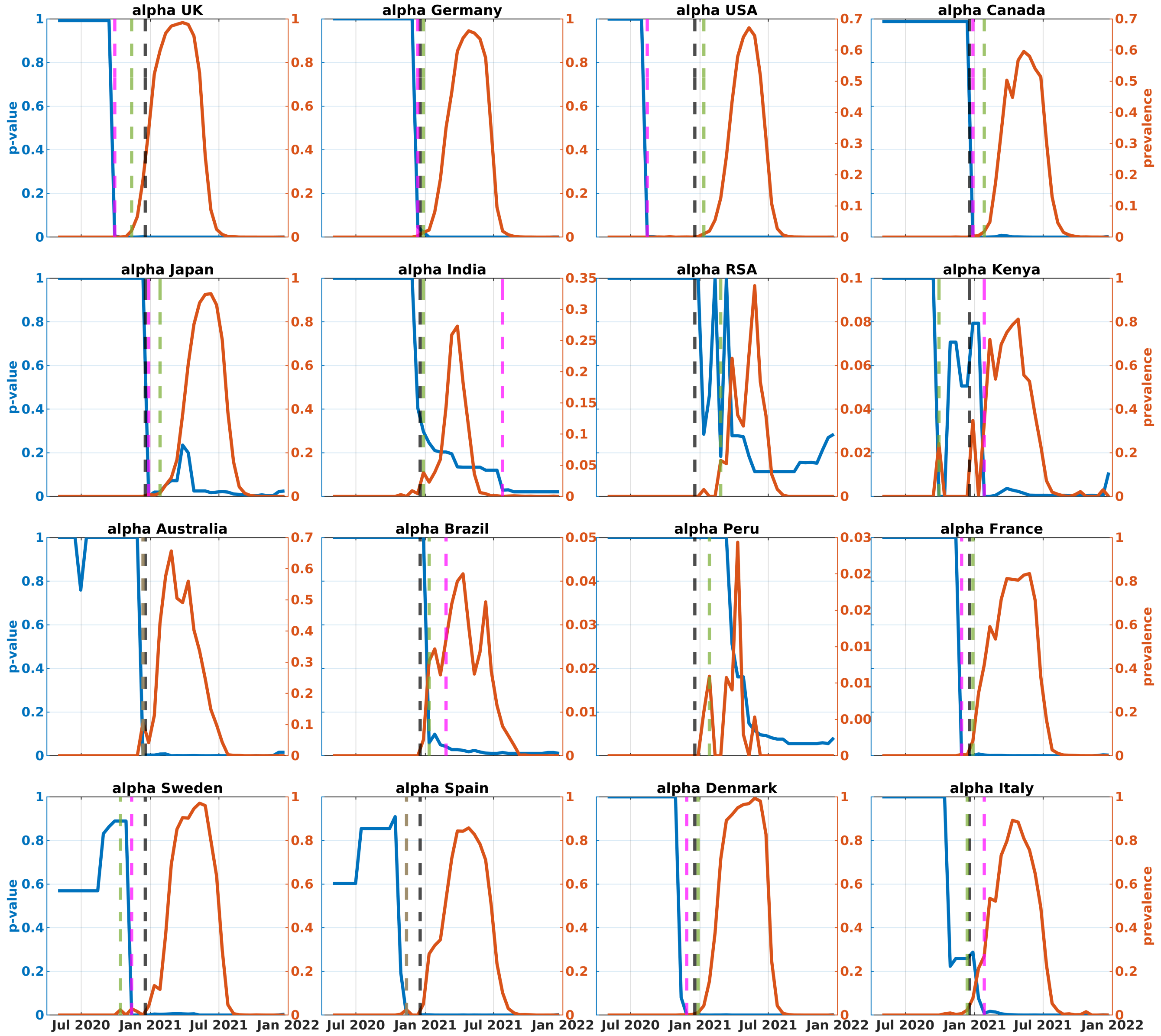


**Fig. A22**: $p$-values (blue) and prevalences (red) of Alpha variant in the analyzed countries (second truncated dataset). Black, green, and magenta lines represent the times of VOC designation, achieving 1% prevalence, and becoming significantly dense, respectively.

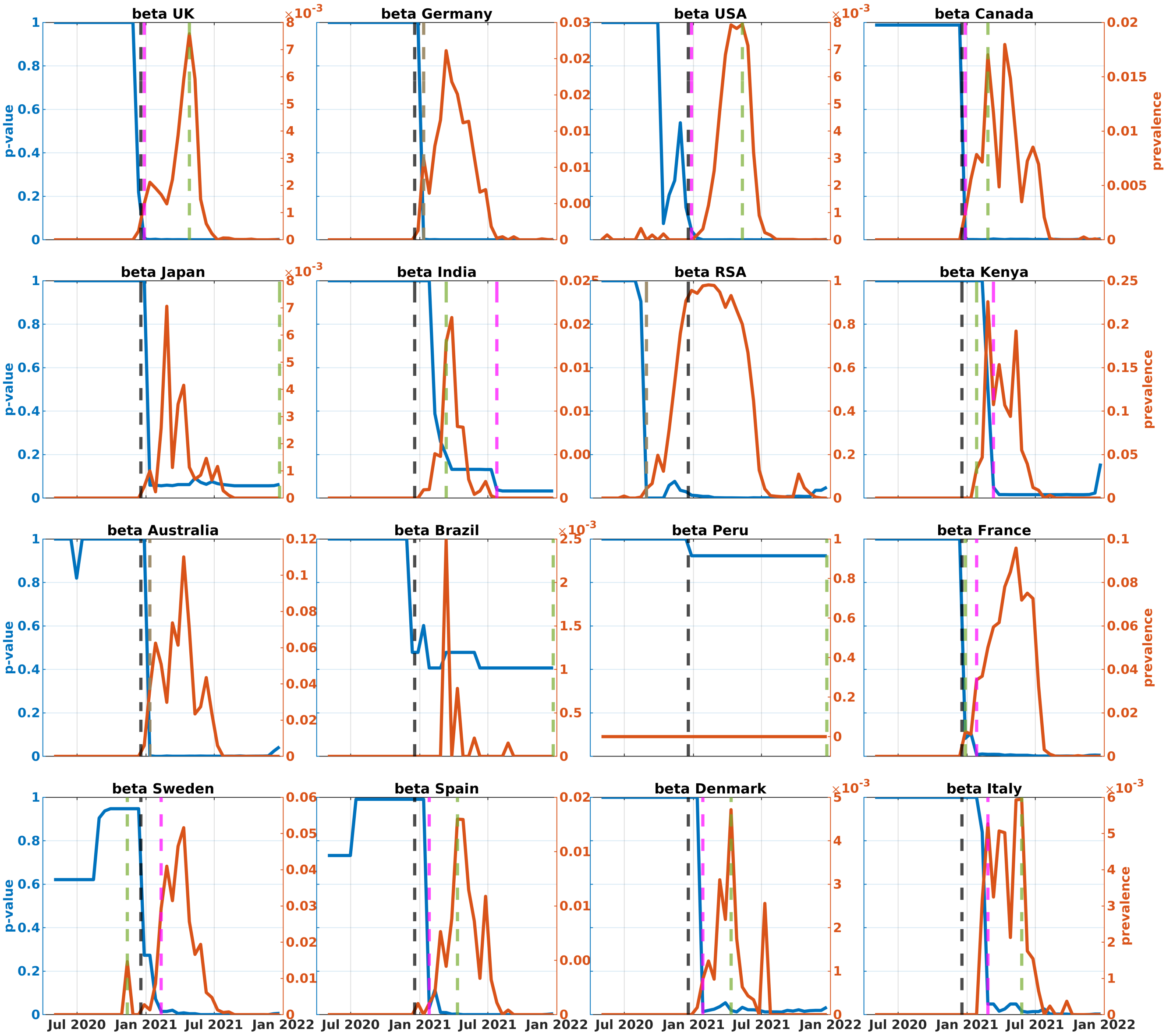


**Fig. A23**: $p$-values (blue) and prevalences (red) of Beta variant in the analyzed countries (second truncated dataset). Black, green, and magenta lines represent the times of VOC designation, achieving 1% prevalence, and becoming significantly dense, respectively.

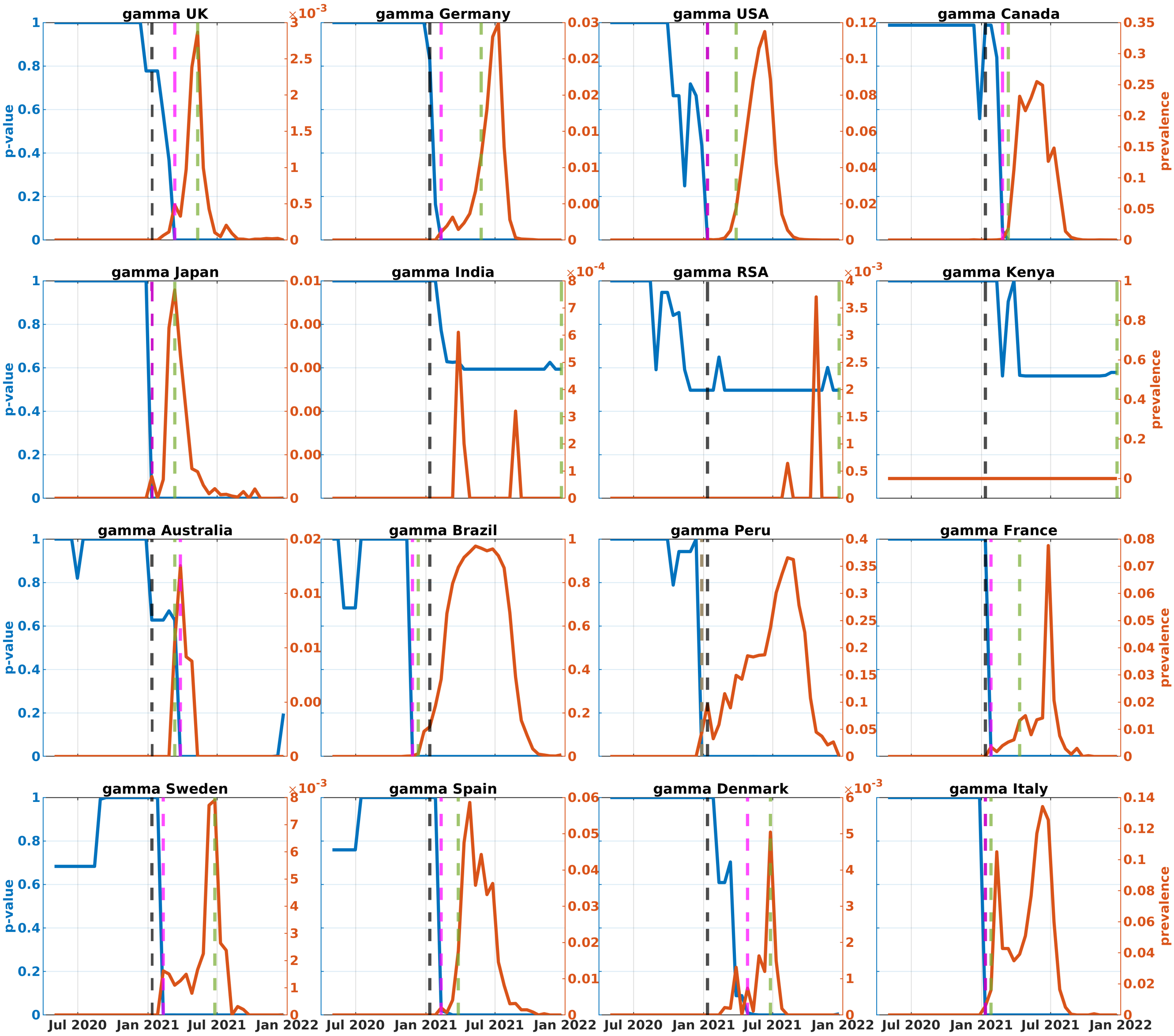


**Fig. A24**: $p$-values (blue) and prevalences (red) of Gamma variant in the analyzed countries (second truncated dataset). Black, green, and magenta lines represent the times of VOC designation, achieving 1% prevalence, and becoming significantly dense, respectively.

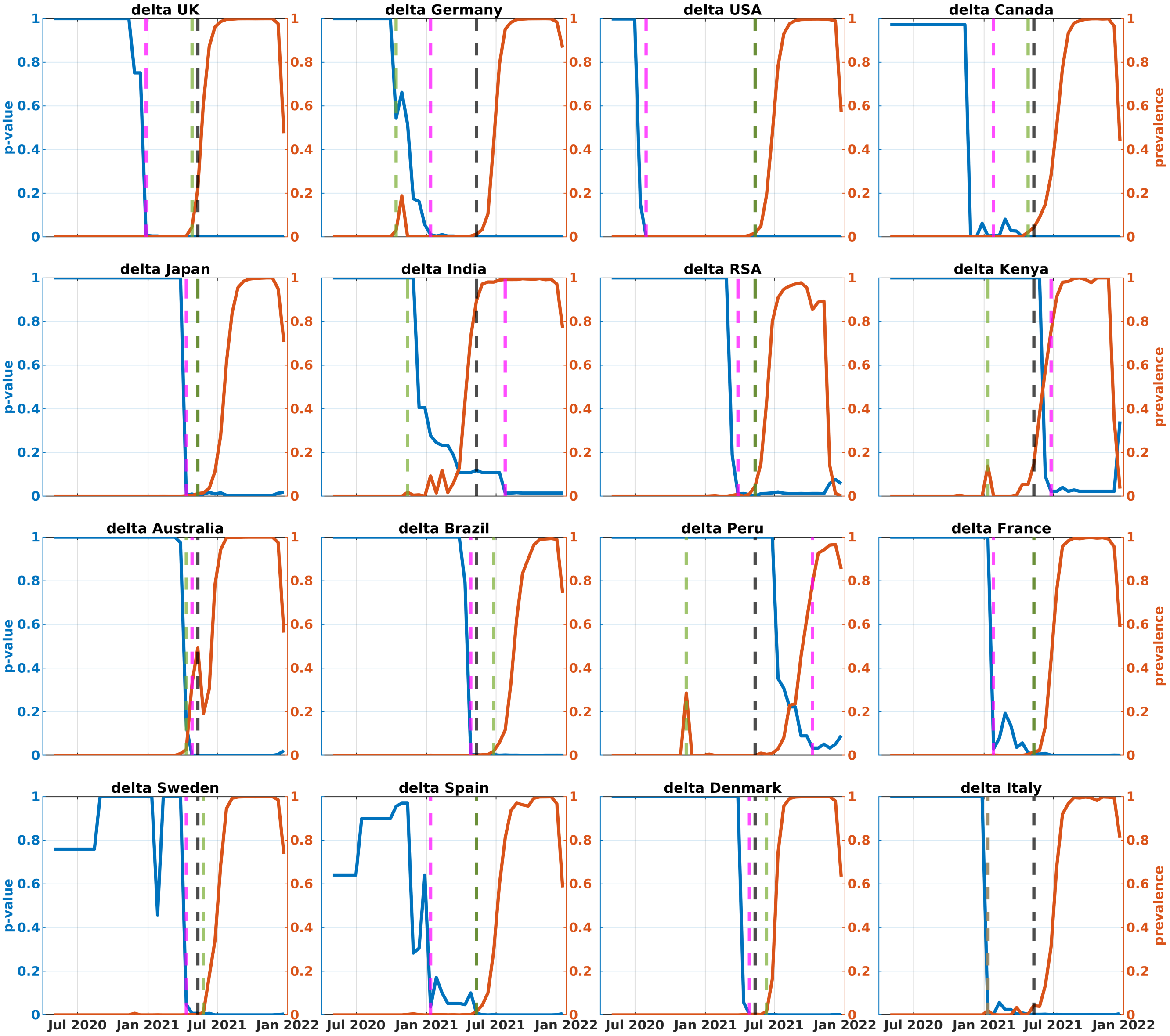


**Fig. A25**: $p$-values (blue) and prevalences (red) of Delta variant in the analyzed countries (second truncated dataset). Black, green, and magenta lines represent the times of VOC designation, achieving 1% prevalence, and becoming significantly dense, respectively.

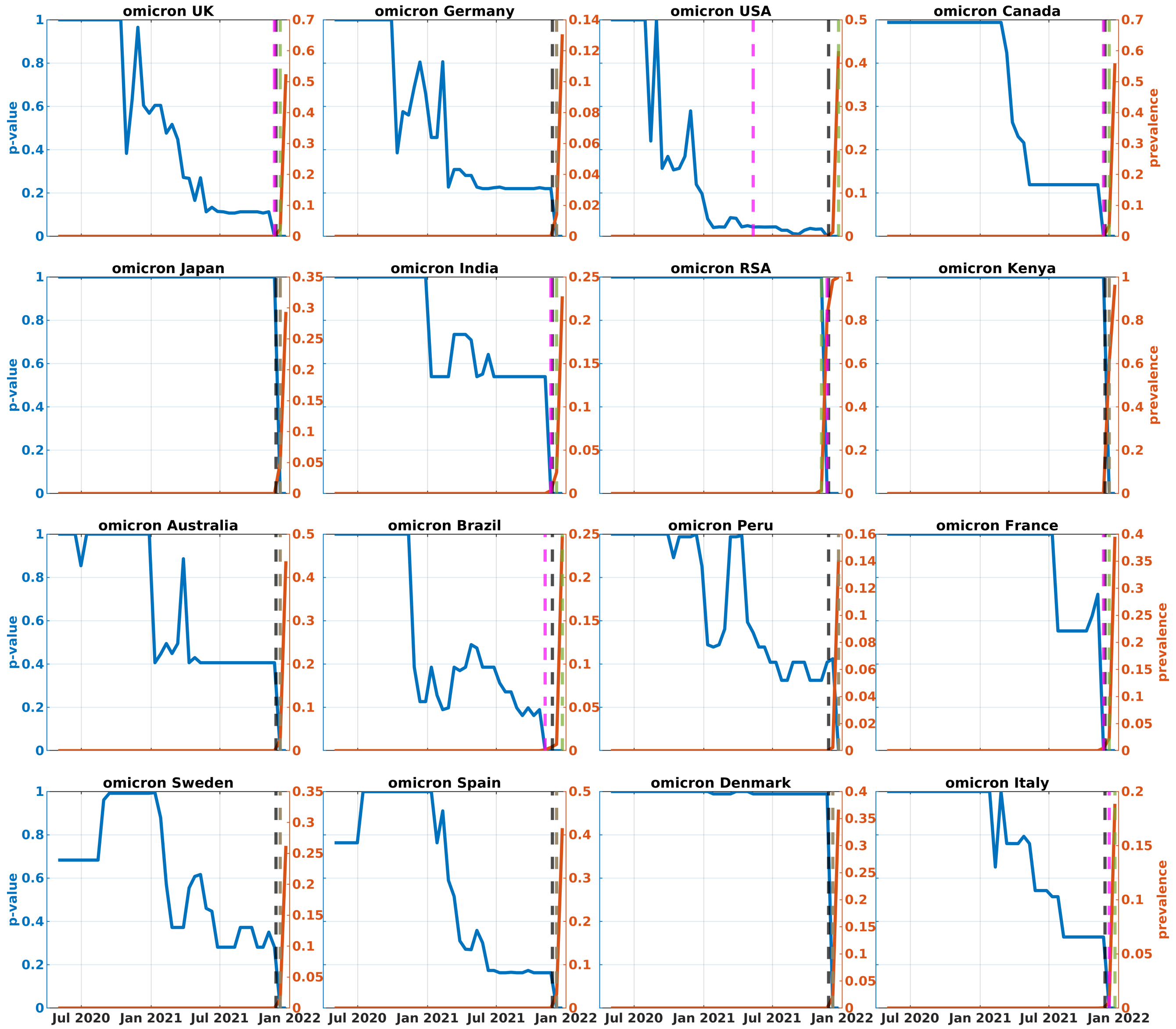


**Fig. A26**: $p$-values (blue) and prevalences (red) of Omicron variant in the analyzed countries (second truncated dataset). Black, green, and magenta lines represent the times of VOC designation, achieving 1% prevalence, and becoming significantly dense, respectively.

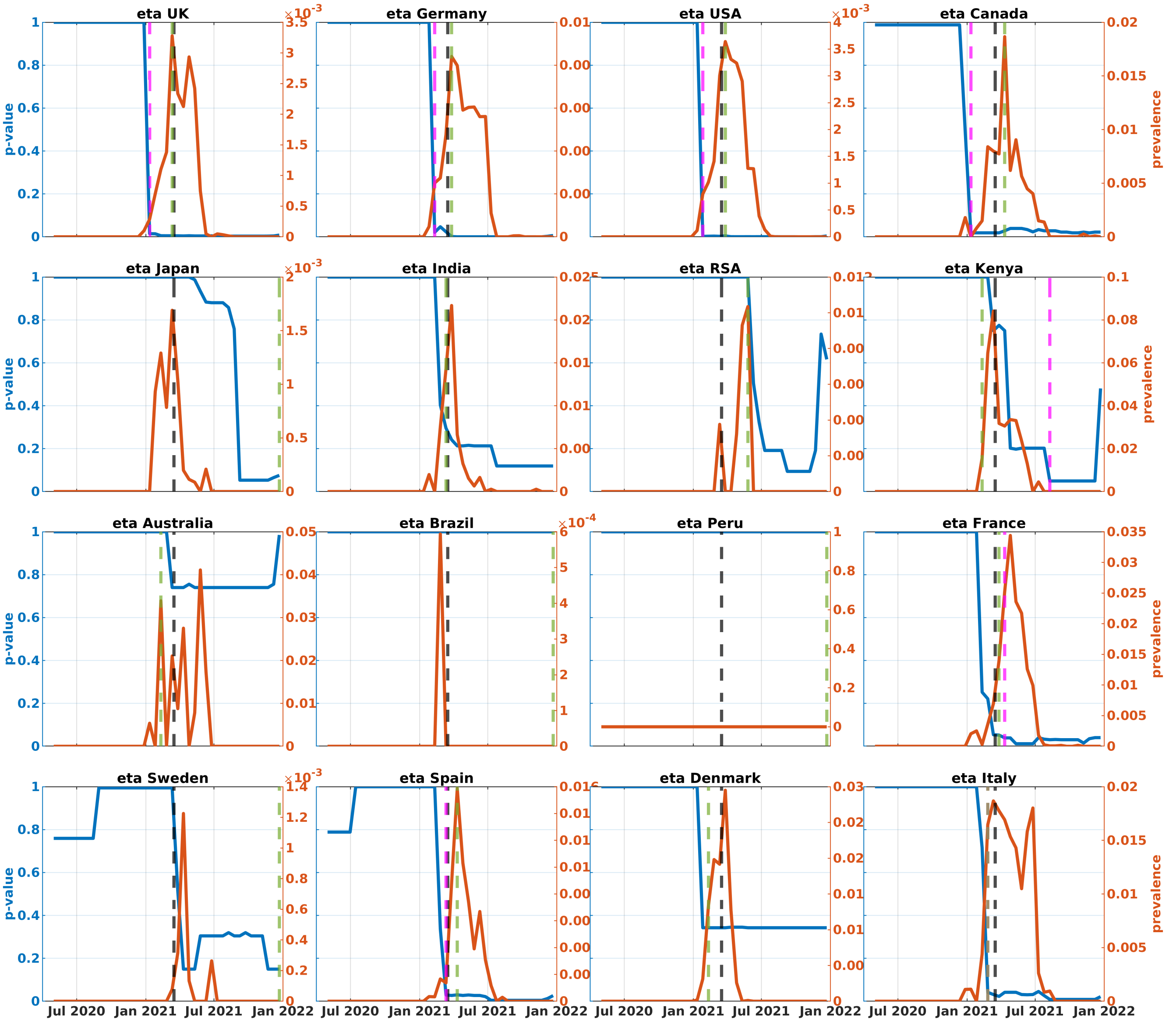


**Fig. A27**: $p$-values (blue) and prevalences (red) of Eta variant in the analyzed countries (second truncated dataset). Black, green, and magenta lines represent the times of VOC designation, achieving 1% prevalence, and becoming significantly dense, respectively.

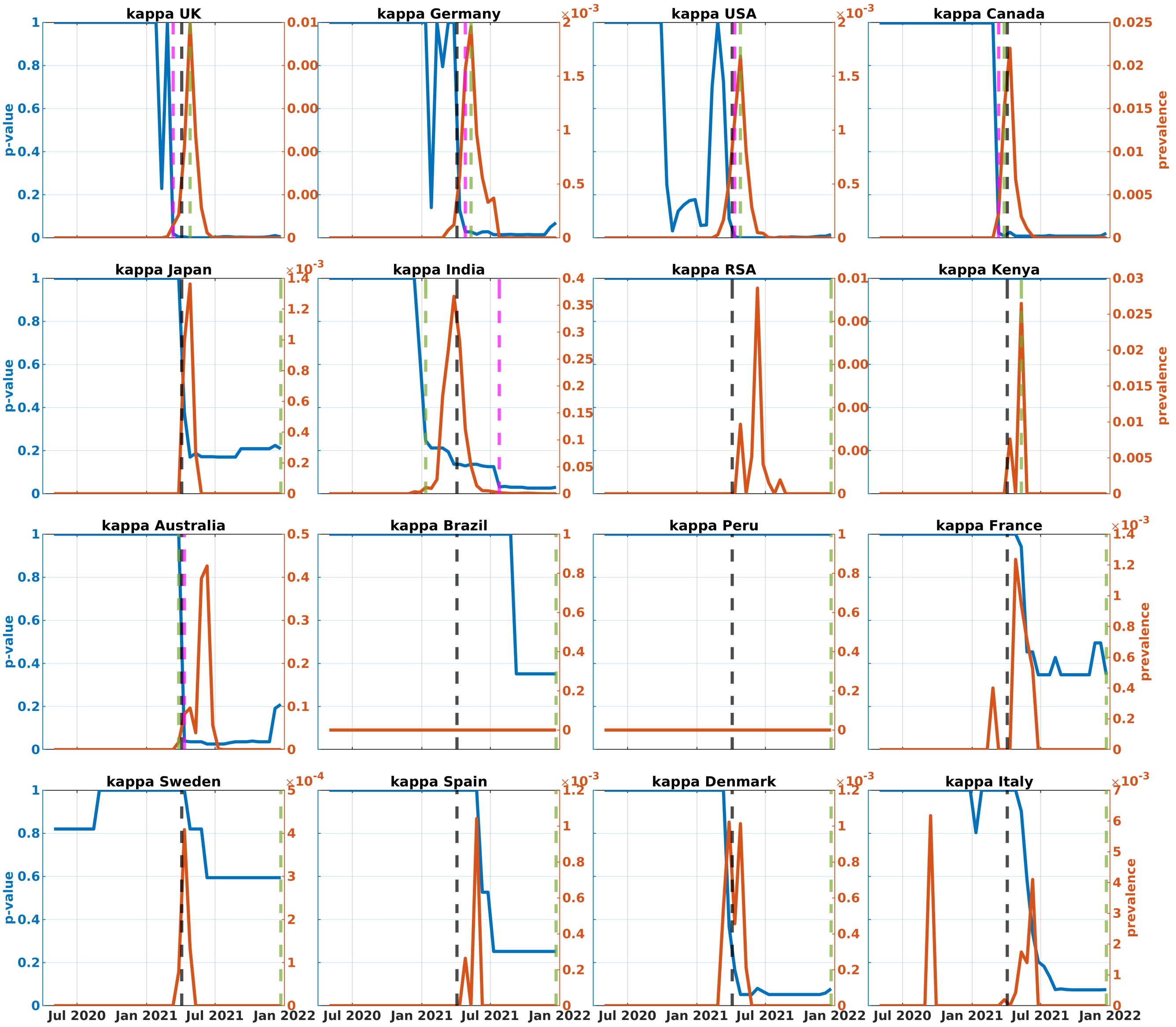


**Fig. A28**: $p$-values (blue) and prevalences (red) of Kappa variant in the analyzed countries (second truncated dataset). Black, green, and magenta lines represent the times of VOC designation, achieving 1% prevalence, and becoming significantly dense, respectively.

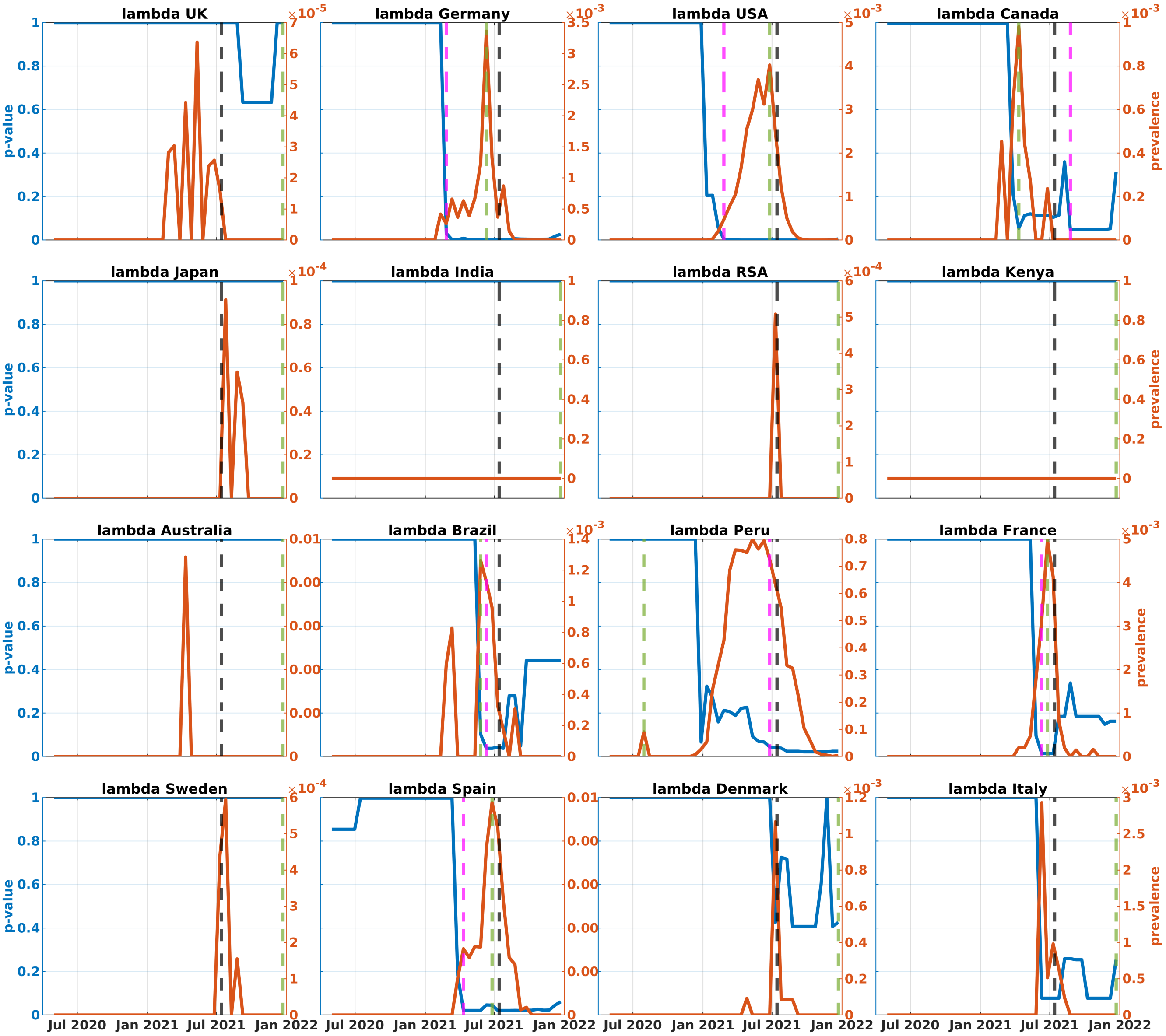


**Fig. A29**: $p$-values (blue) and prevalences (red) of Lambda variant in the analyzed countries (second truncated dataset). Black, green, and magenta lines represent the times of VOC designation, achieving 1% prevalence, and becoming significantly dense, respectively.

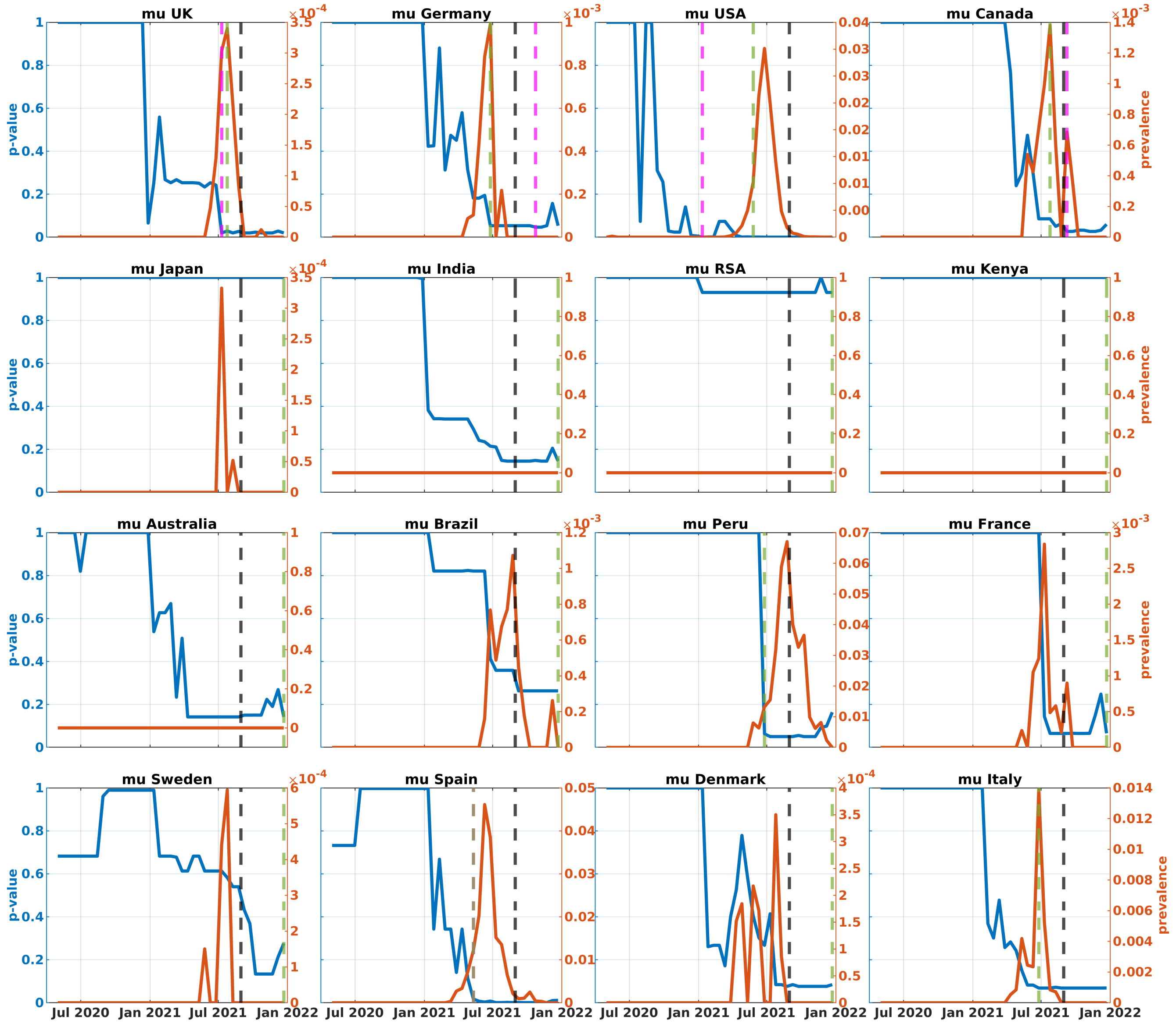


**Fig. A30**: $p$-values (blue) and prevalences (red) of Mu variant in the analyzed countries (second truncated dataset). Black, green, and magenta lines represent the times of VOC designation, achieving 1% prevalence, and becoming significantly dense, respectively.

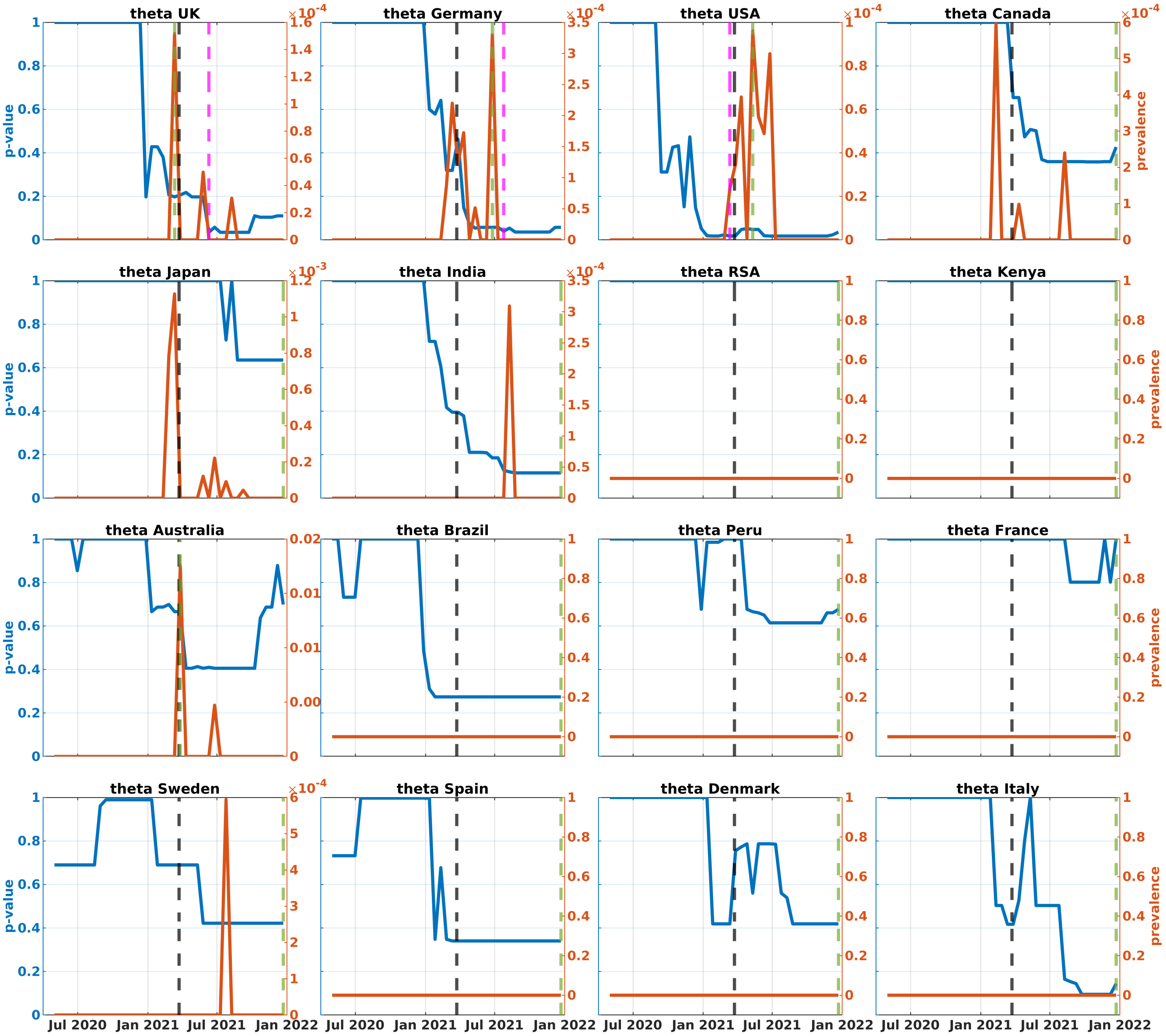


**Fig. A31**: $p$-values (blue) and prevalences (red) of Theta variant in the analyzed countries (second truncated dataset). Black, green, and magenta lines represent the times of VOC designation, achieving 1% prevalence, and becoming significantly dense, respectively.

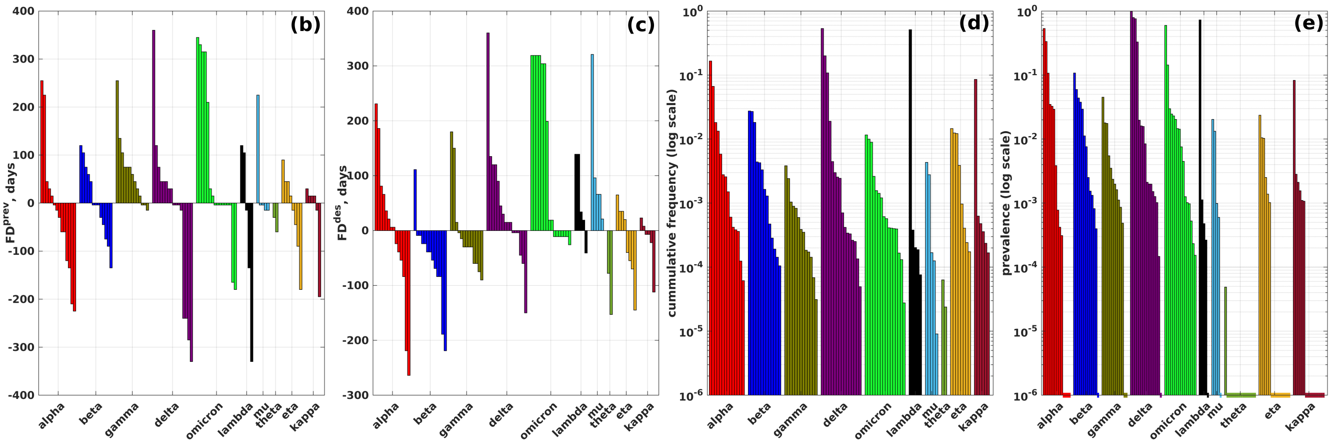


**Fig. A32**: **Density-based adjusted $p$-values of VOCs/VOIs (complete dataset).** (b) and (c): Forecasting depths (y-axis) in relation to the 1% prevalence time and WHO designation time for each analyzed VOC/VOI across different countries. (d) and (e): Cumulative frequencies and prevalences for VOCs/VOIs across various countries at the times when they become significantly dense (in a logarithmic scale). Dashed lines at the bottom of the plot indicate that the variants reached significant density at frequencies/prevalences of 0.

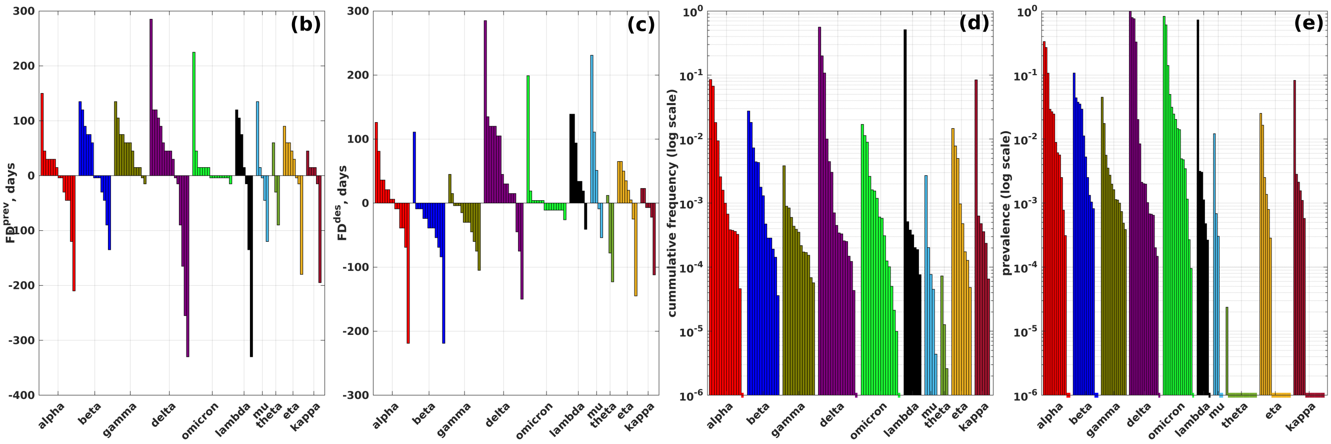


**Fig. A33**: **Density-based adjusted $p$-values of VOCs/VOIs (second truncated dataset).** (b) and (c): Forecasting depths (y-axis) in relation to the 1% prevalence time and WHO designation time for each analyzed VOC/VOI across different countries. (d) and (e): Cumulative frequencies and prevalences for VOCs/VOIs across various countries at the times when they become significantly dense (in a logarithmic scale). Dashed lines at the bottom of the plot indicate that the variants reached significant density at frequencies/prevalences of 0.

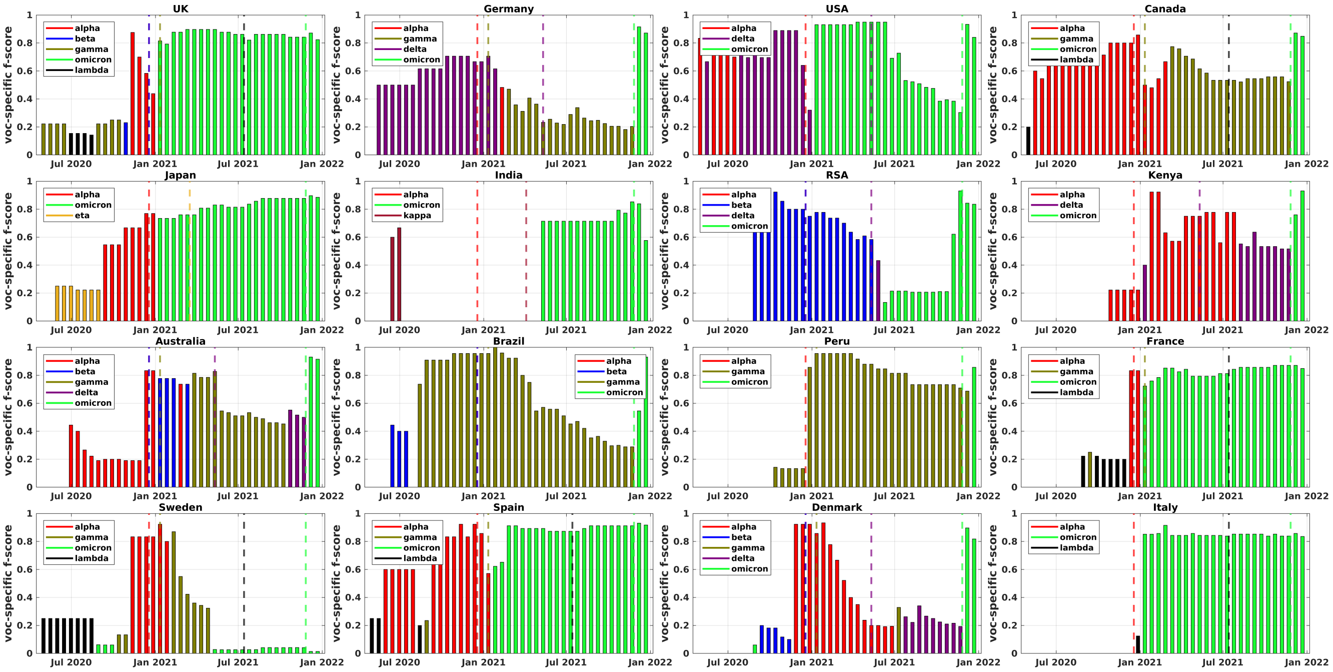


**Fig. A34**: Comparison between VOCs and densest subnetworks of temporal epistatic networks for individual countries (complete dataset). At each time point, bar color code corresponds to the VOC closest to the inferred densest subnetwork, and the bar hight is equal to the respective $f$-score. Colored dashed lines mark times when specific VOCs were designated by WHO.

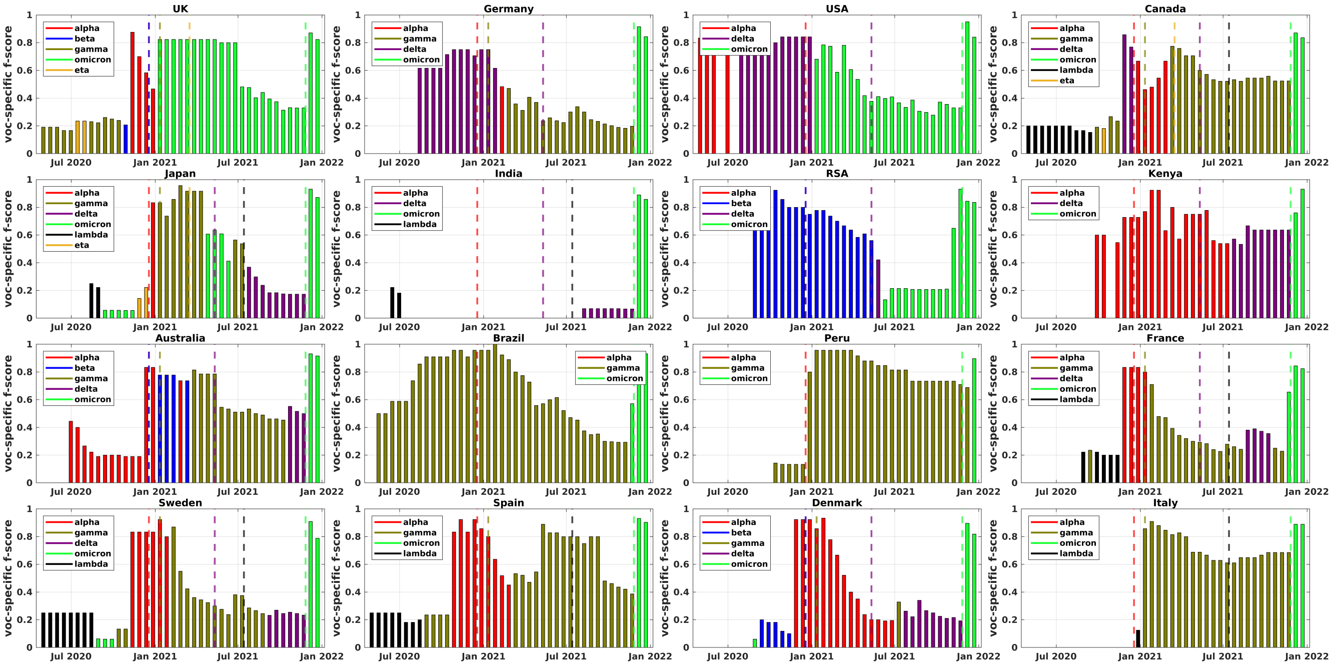

**Fig. A35**: Comparison between VOCs and densest subnetworks of temporal epistatic networks for individual countries (first truncated dataset). At each time point, bar color code corresponds to the VOC closest to the inferred densest subnetwork, and the bar hight is equal to the respective $f$-score. Colored dashed lines mark times when specific VOCs were designated by WHO.

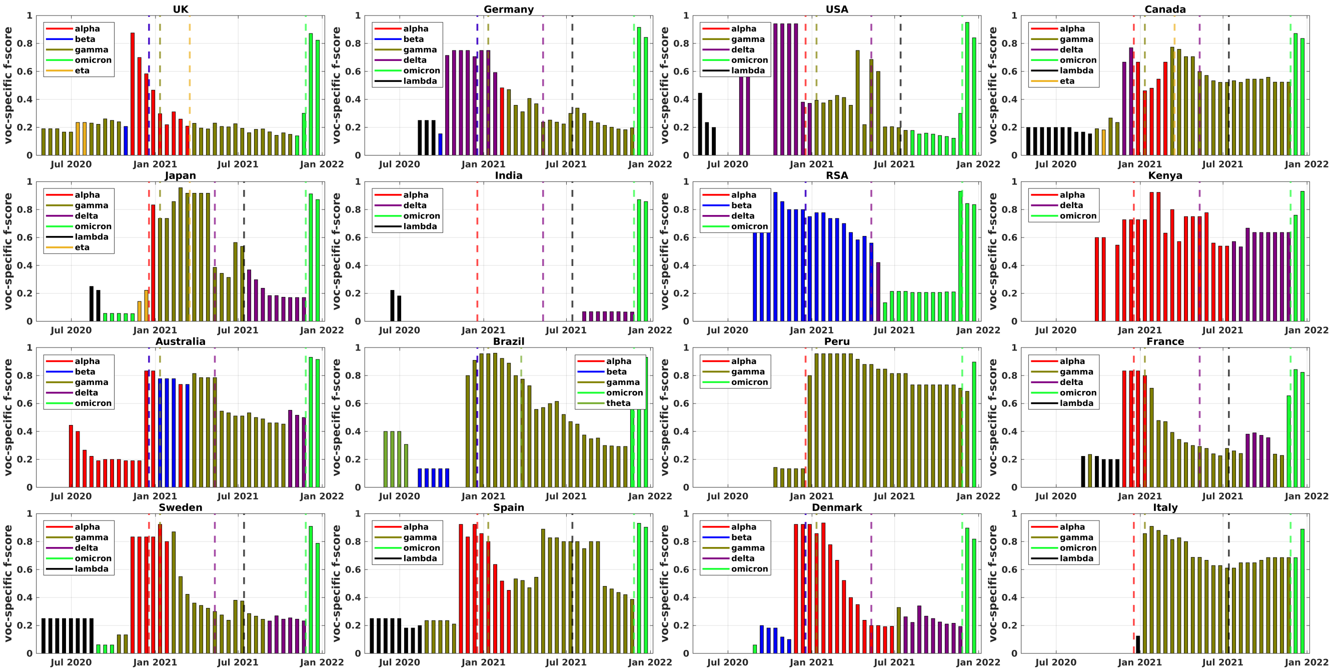


**Fig. A36**: Comparison between VOCs and densest subnetworks of temporal epistatic networks for individual countries (second truncated dataset). At each time point, bar color code corresponds to the VOC closest to the inferred densest subnetwork, and the bar hight is equal to the respective $f$-score. Colored dashed lines mark times when specific VOCs were designated by WHO.

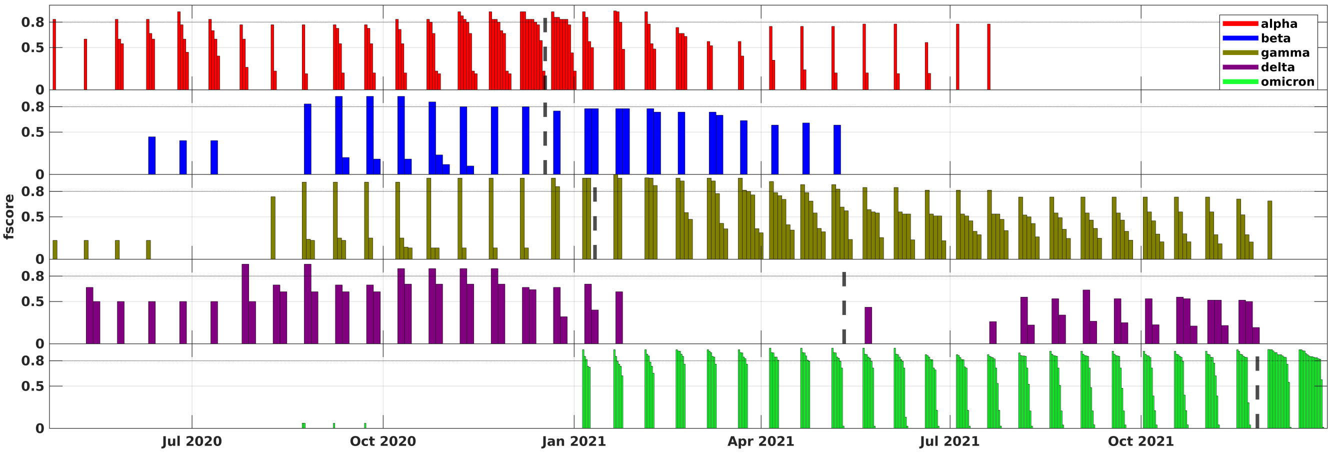


**Fig. A37**: **Comparison of the densest subnetworks from temporal coordinated substitution networks (aggregated over 16 countries) with VOCs for the complete dataset.** Each bar in the plot represents a specific VOC. For every time point, the bars display the densest subgraphs from different countries that are most similar to that VOC, with the height of the bars indicating the corresponding $f$-scores. Colored dashed lines highlight the moments when the WHO designated the VOCs.

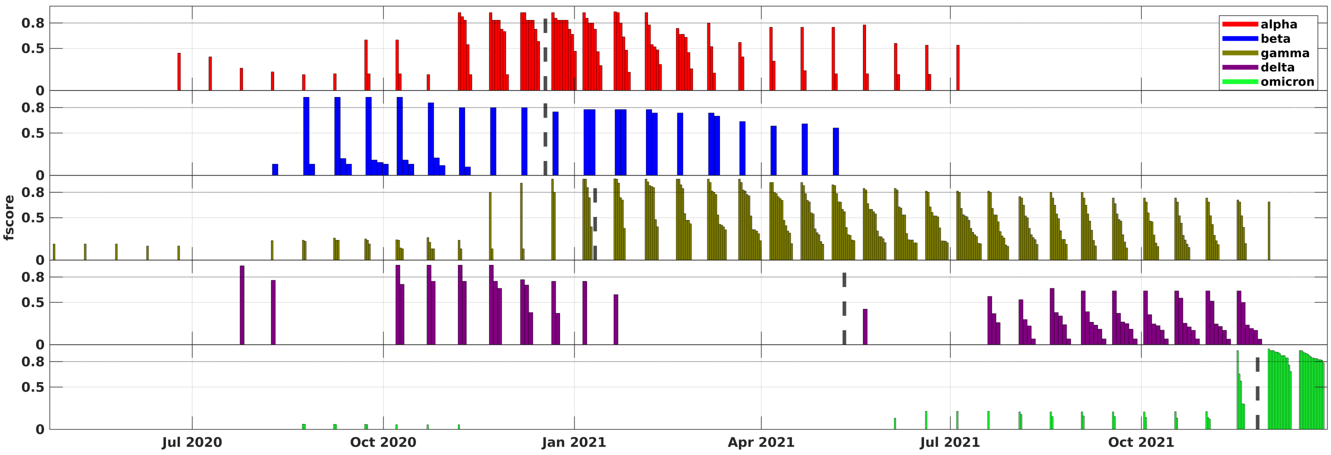


**Fig. A38**: **Comparison of the densest subnetworks from temporal coordinated substitution networks (aggregated over 16 countries) with VOCs for the second truncated dataset.** Each bar in the plot represents a specific VOC. For every time point, the bars display the densest subgraphs from different countries that are most similar to that VOC, with the height of the bars indicating the corresponding $f$-scores. Colored dashed lines highlight the moments when the WHO designated the VOCs.

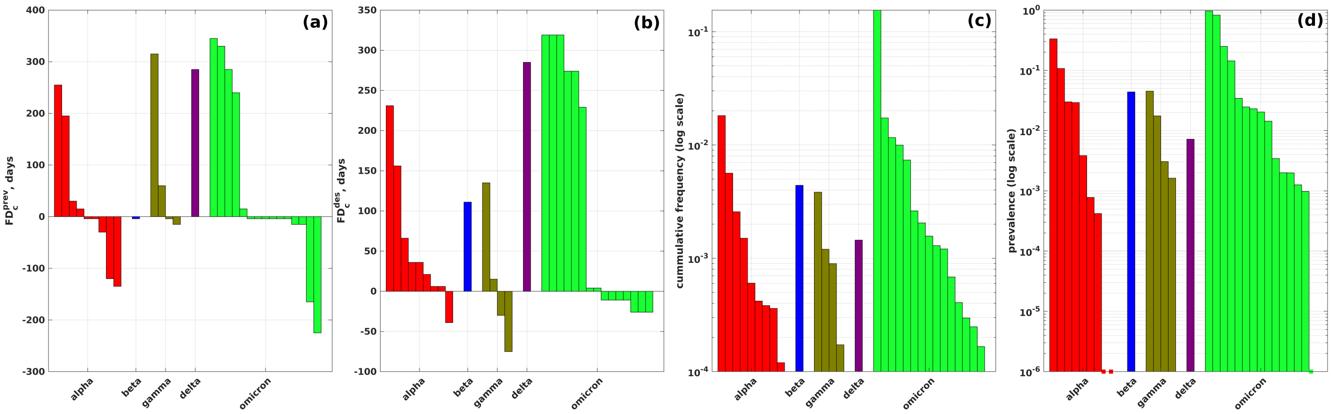


**Fig. A39**: **Summary of comparison between VOCs and densest subnetworks of temporal epistatic networks for all countries (complete dataset)**. (a) and (b): forecasting depths (y-axis) with respect to the 1% prevalence time and WHO designation time for each analyzed VOCs over different countries. (c) and (d): cumulative frequencies and prevalences of VOCs over different countries at earliest times when they are at least 80% identical to densest subgraphs of epistatic networks (in logarithmic scale).

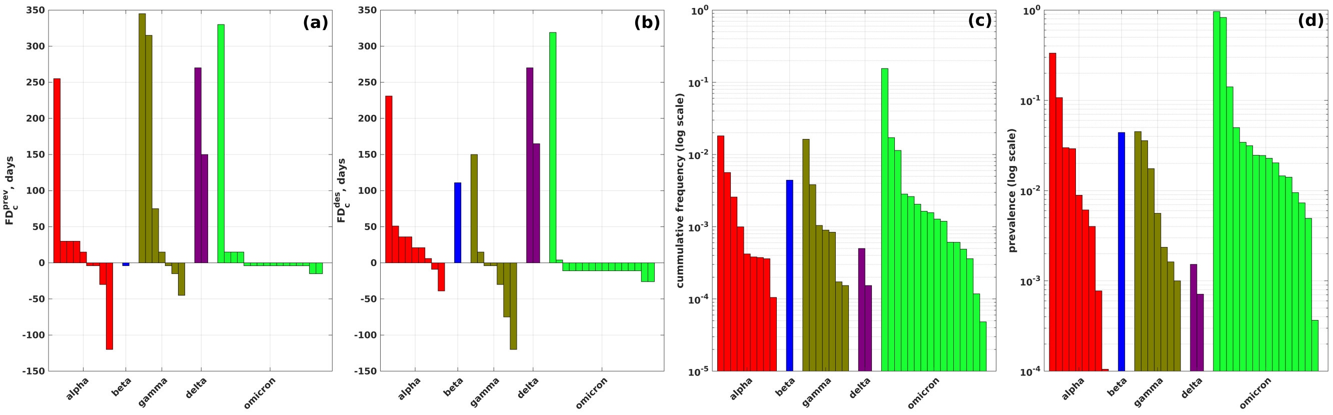


**Fig. A40**: **Summary of comparison between VOCs and densest subnetworks of temporal epistatic networks for all countries (first truncated dataset)**. (a) and (b): forecasting depths (y-axis) with respect to the 1% prevalence time and WHO designation time for each analyzed VOCs over different countries. (c) and (d): cumulative frequencies and prevalences of VOCs over different countries at earliest times when they are at least 80% identical to densest subgraphs of epistatic networks (in logarithmic scale).

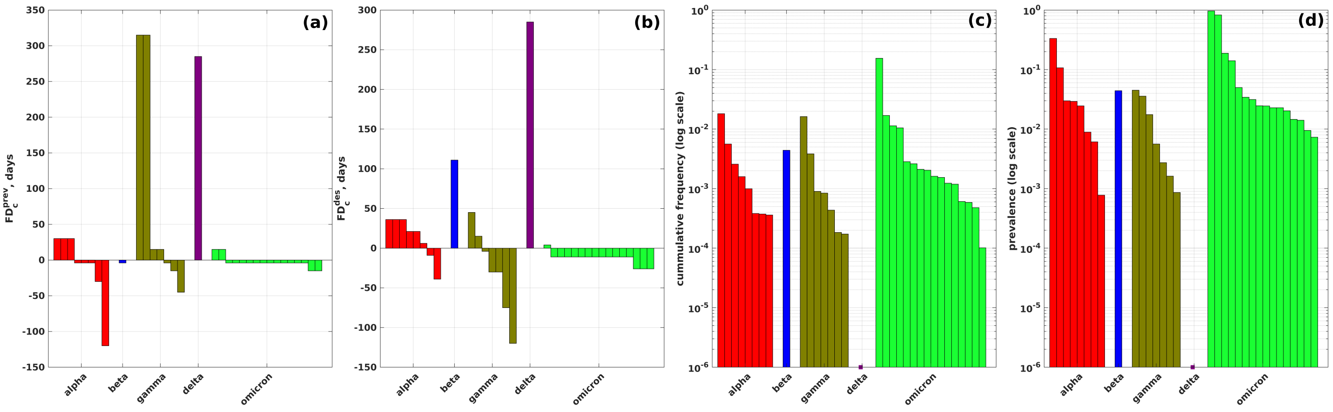


**Fig. A41**: **Summary of comparison between VOCs and densest subnetworks of temporal epistatic networks for all countries (second truncated dataset)**. (a) and (b): forecasting depths (y-axis) with respect to the 1% prevalence time and WHO designation time for each analyzed VOCs over different countries. (c) and (d): cumulative frequencies and prevalences of VOCs over different countries at earliest times when they are at least 80% identical to densest subgraphs of epistatic networks (in logarithmic scale).

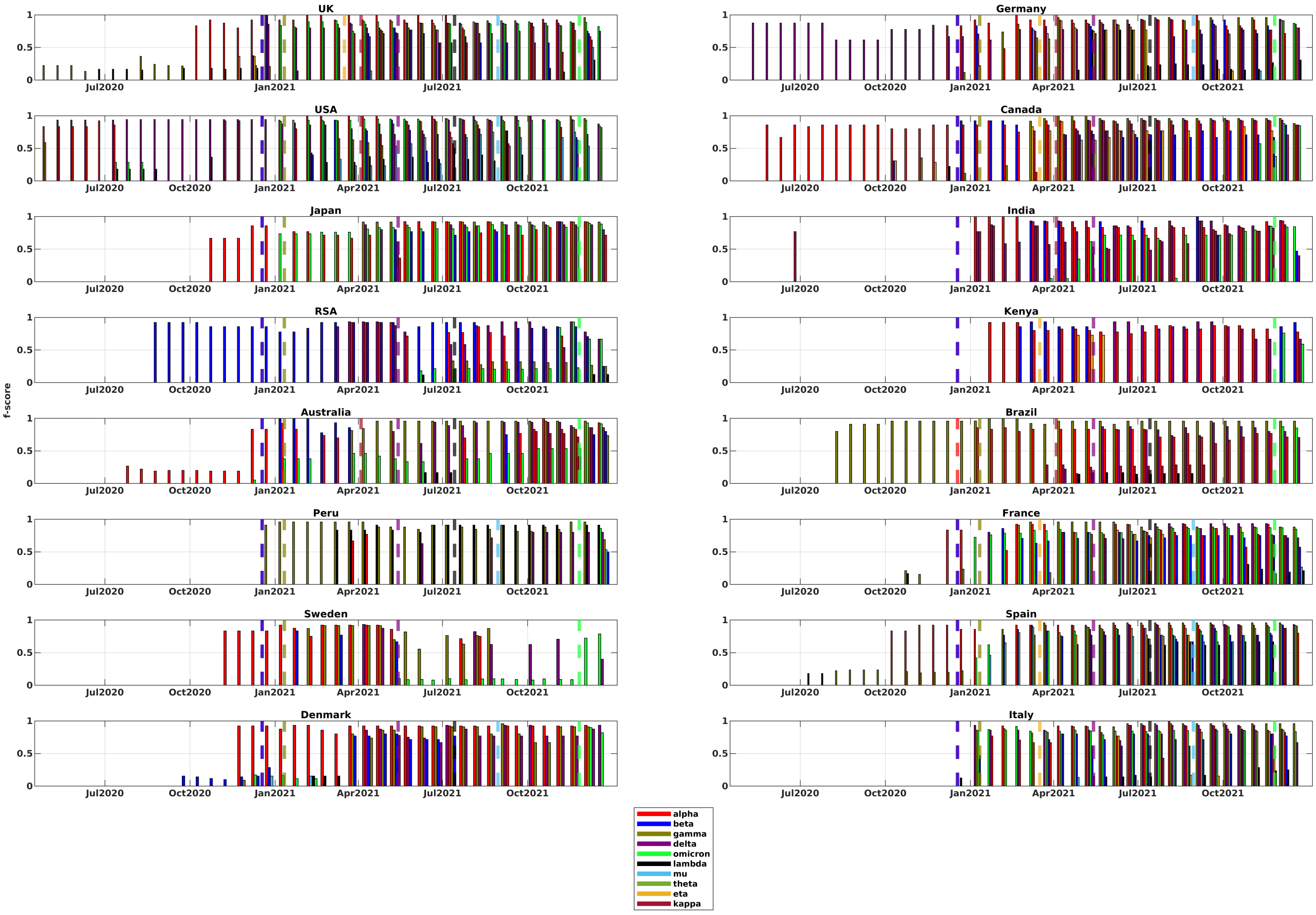


**Fig. A42**: **Comparison between VOCs and inferred haplotypes for individual countries** (complete dataset). At each time point, each bar represents an inferred haplotype closest to a particular VOC, the bar height is equal to the respective $f$-score. Colored dashed lines mark times when specific VOCs/VOIs were designated by WHO.

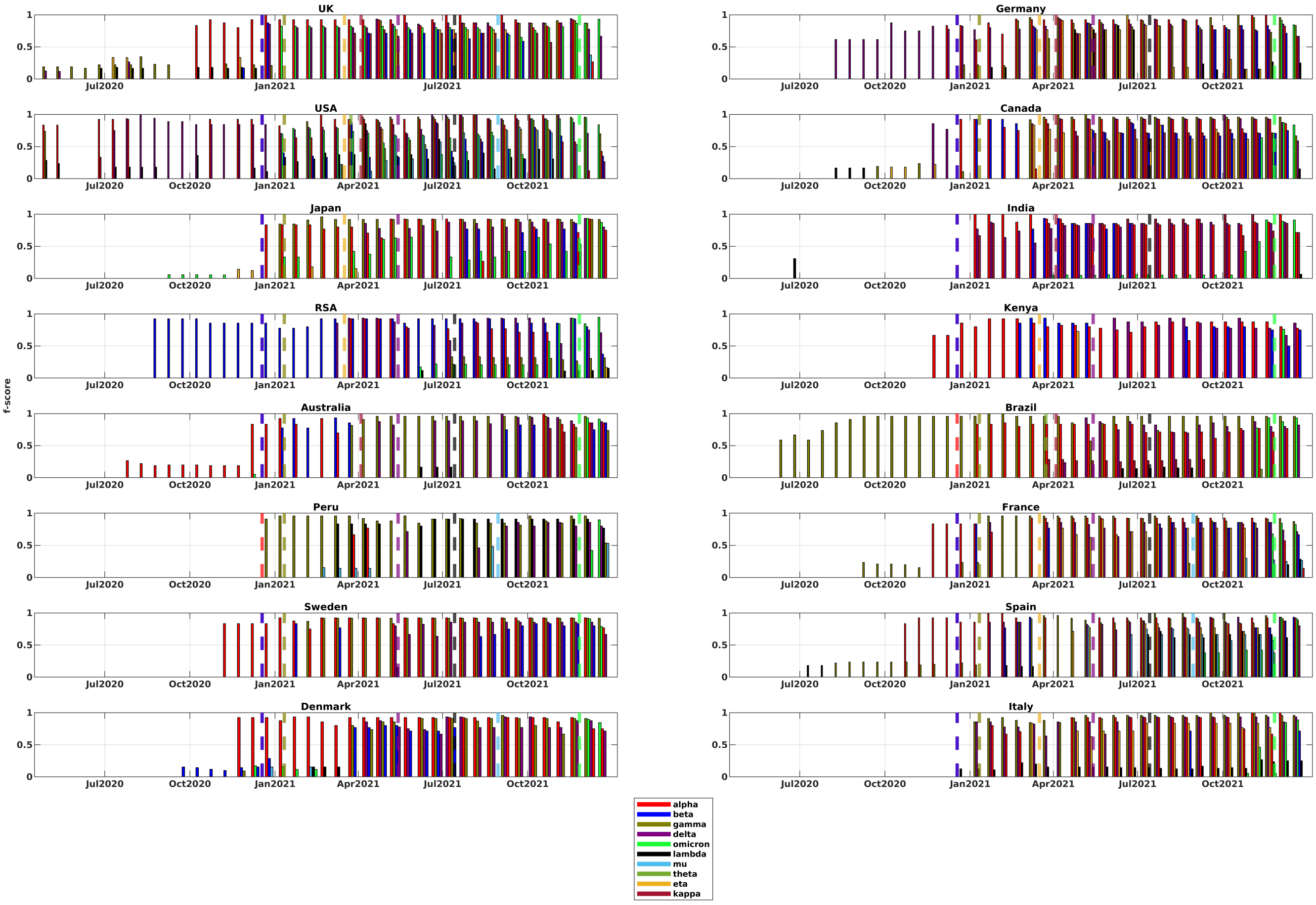


**Fig. A43**: **Comparison between VOCs and inferred haplotypes for individual countries** (complete dataset). At each time point, each bar represents an inferred haplotype closest to a particular VOC, the bar height is equal to the respective $f$-score. Colored dashed lines mark times when specific VOCs/VOIs were designated by WHO.

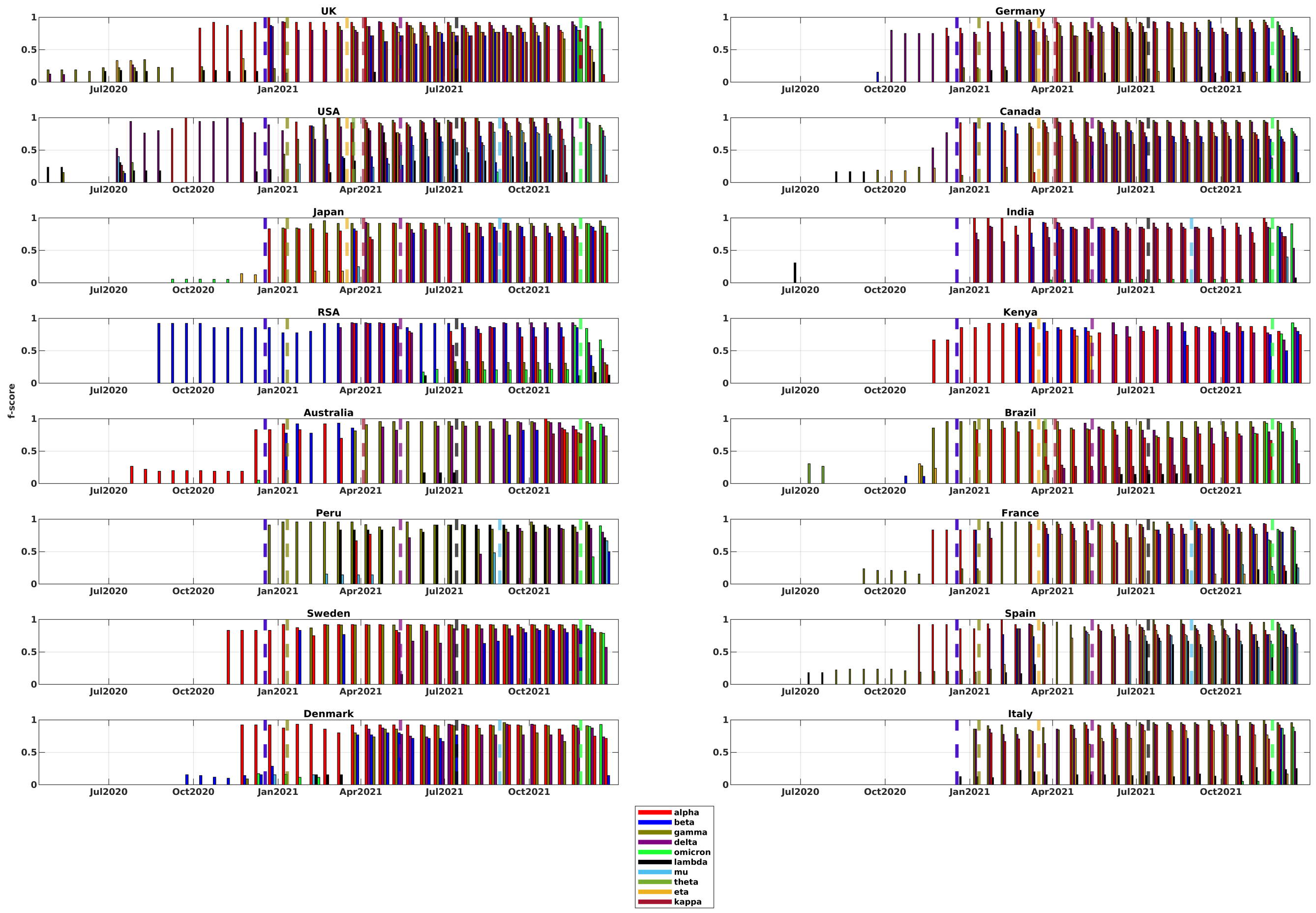


**Fig. A44**: **Comparison between VOCs and inferred haplotypes for individual countries** (complete dataset). At each time point, each bar represents an inferred haplotype closest to a particular VOC, the bar height is equal to the respective $f$-score. Colored dashed lines mark times when specific VOCs/VOIs were designated by WHO.

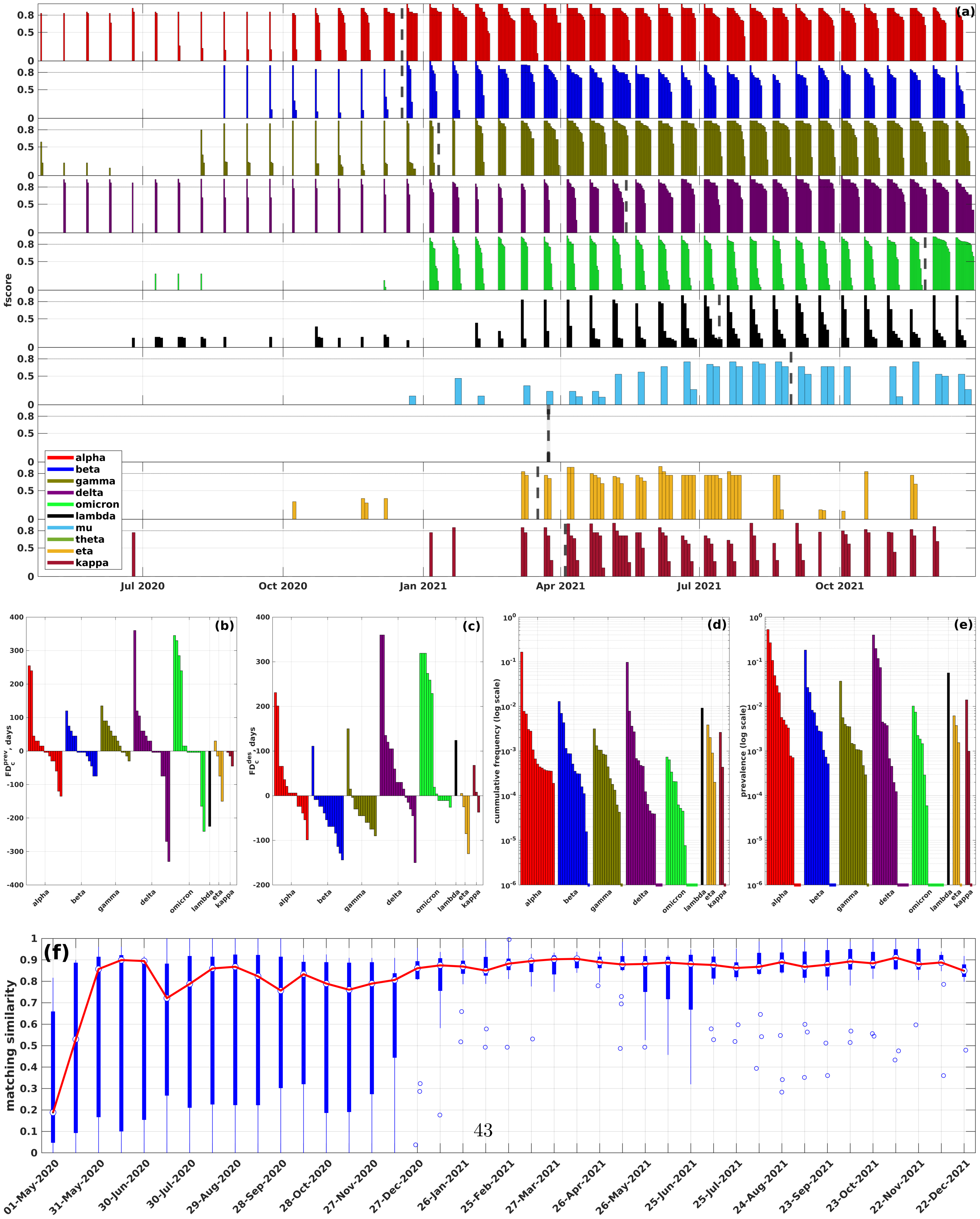




**Fig. A45**: **(a) Summary of comparison between VOCs/VOIs and inferred haplotypes** (complete dataset). Each bar plot depicts the comparison results for a particular VOC/VOI; at each time point, bars correspond to inferred haplotypes from different countries closest to that VOC, and the bar heights are equal to the respective $f$-scores. Colored dashed lines mark times when the VOCs were designated by WHO. **(b) and (c): forecasting depths** (y-axis) with respect to the 1% prevalence time and WHO designation time for each analyzed VOCs/VOIs over different countries. **(d) and (e): cumulative frequencies and prevalences** of VOCs/VOIs over different countries at first variant call times (in logarithmic scale). Dashed lines at the bottom of the plot signify that the corresponding variants were detected at cumulative frequencies or prevalences 0. **(f) Precision of haplotype inference.** Blue box plot depicts summary statistics of matching similarity of $n = 16$ countries over $T = 21$ time points. The bottom and top of each box are the 25th and 75th percentiles, whiskers represent minimum and maximum values, white dot is a median. Red plot depicts the dynamics of median matching similarity over time.

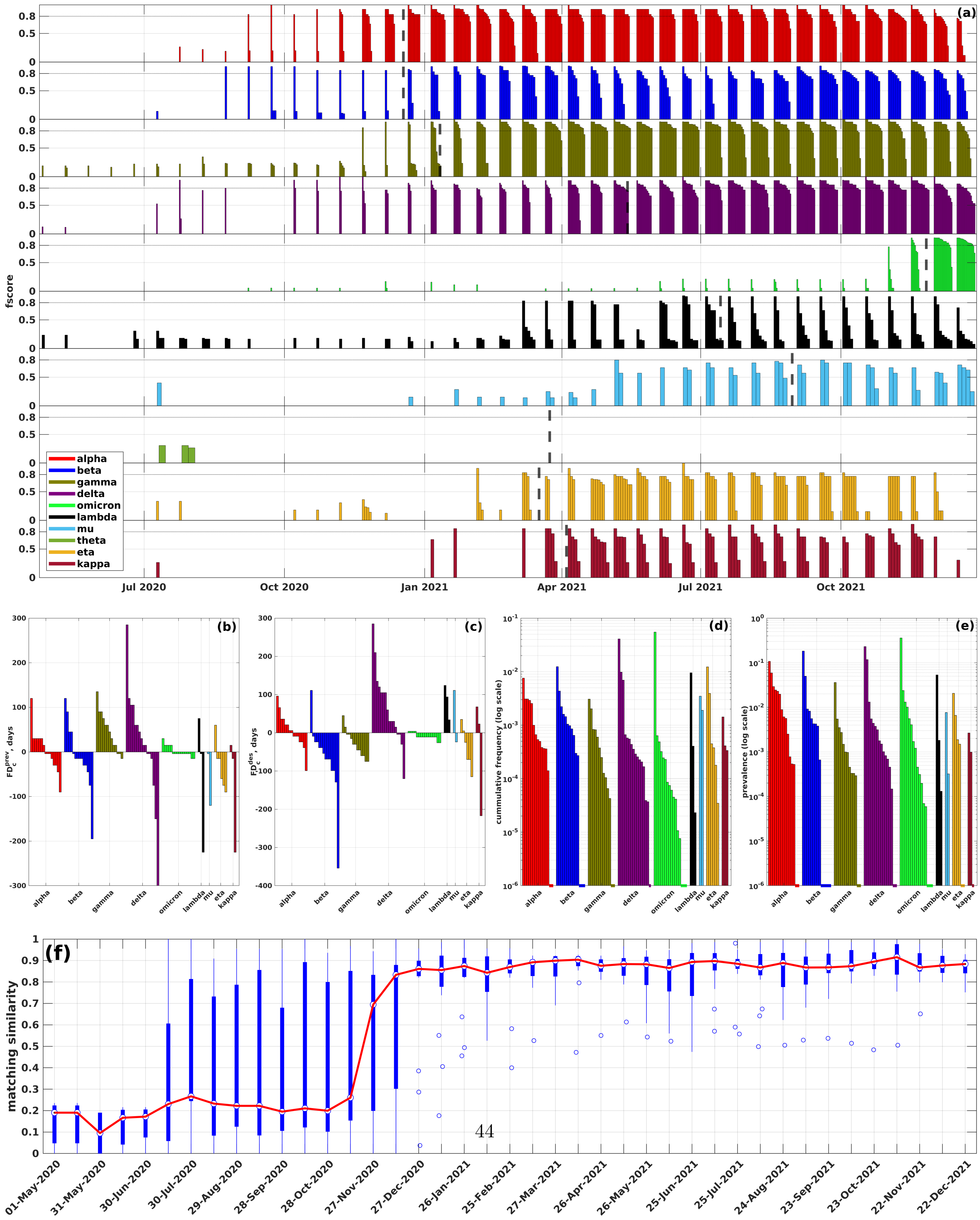




**Fig. A46**: **(a) Summary of comparison between VOCs/VOIs and inferred haplotypes** (complete dataset). Each bar plot depicts the comparison results for a particular VOC/VOI; at each time point, bars correspond to inferred haplotypes from different countries closest to that VOC, and the bar heights are equal to the respective $f$-scores. Colored dashed lines mark times when the VOCs were designated by WHO. **(b) and (c): forecasting depths** (y-axis) with respect to the 1% prevalence time and WHO designation time for each analyzed VOCs/VOIs over different countries. **(d) and (e): cumulative frequencies and prevalences** of VOCs/VOIs over different countries at first variant call times (in logarithmic scale). Dashed lines at the bottom of the plot signify that the corresponding variants were detected at cumulative frequencies or prevalences 0. **(f) Precision of haplotype inference.** Blue box plot depicts summary statistics of matching similarity of $n = 16$ countries over $T = 21$ time points. The bottom and top of each box are the 25th and 75th percentiles, whiskers represent minimum and maximum values, white dot is a median. Red plot depicts the dynamics of median matching similarity over time.
